# EML4-ALK variant-specific genetic interactions shape lung tumorigenesis

**DOI:** 10.1101/2024.08.26.609730

**Authors:** Alberto Diaz-Jimenez, Emily G. Shuldiner, Kalman Somogyi, Oscar Gonzalez, Filiz Akkas, Christopher W. Murray, Laura Andrejka, Min K. Tsai, Benedikt Brors, Smruthy Sivakumar, Saumya D. Sisoudiya, Ethan S. Sokol, Dmitri A. Petrov, Monte M. Winslow, Rocio Sotillo

## Abstract

Oncogenic fusions of EML4 and ALK occur in ∼5% of lung adenocarcinomas. More than 15 EML4-ALK variants with distinct breakpoints within *EML4* have been identified, but the functional differences between these variants remain poorly understood. Here we use CRISPR/Cas9 somatic genome editing to generate autochthonous mouse models of the two most common EML4-ALK variants, V1 and V3, and show that V3 is more oncogenic than V1. By integrating these models with multiplexed genome editing, we quantify the effects of 29 putative tumor suppressor genes on V1- and V3-driven lung cancer growth *in vivo* and show that many tumor suppressor genes have dramatically variant-specific effects on tumorigenesis. Analysis of a novel dataset representing the largest human EML4-ALK lung cancer cohort to date identified alterations in the genomic landscape depending on the EML4-ALK variant. These findings demonstrate functional heterogeneity among EML4-ALK variants, suggesting that EML4-ALK variants behave more like distinct oncogenes than a uniform entity. More broadly, these findings highlight the dramatic impact of oncogenic fusions partner proteins on tumor biology.

## INTRODUCTION

Cancer genomic studies have uncovered diverse oncogenic chromosomal rearrangements (Sanchez-Vega et al. 2018; Gao et al. 2018; Lee et al. 2019; Consortium et al.). Among these, fusions between echinoderm microtubule-associated protein-like 4 (EML4) and the anaplastic lymphoma kinase (ALK) are oncogenic drivers in 3-7% of lung adenocarcinomas, accounting for more than 100,000 cases worldwide each year (Schneider et al. 2023). *EML4-ALK* fusions are generated by chromosomal inversions which fuse the 3’ end of *ALK* with various breakpoints in *EML4*, resulting in fusion proteins that contain the tyrosine kinase domain of ALK but vary substantially in the length of the EML4 partner protein (Soda et al. 2007; Choi et al. 2008; Sabir et al. 2017; Ou et al. 2020). More than 15 recurrent *EML4-ALK* variants have been identified in patients, with variant 1 (V1; exon 13 of *EML4* fused with exon 20 of *ALK*) and variant 3 (V3; exon 6a/b of *EML4* fused with exon 20 of *ALK*) accounting for approximately 80% of cases (Sabir et al. 2017). *In vitro* studies suggest that there are differences among EML4-ALK variants in stability, localization, and sensitivity to ALK inhibition (Heuckmann et al. 2012; Childress et al. 2018; O’Regan et al. 2020); however, EML4-ALK-driven tumors are typically viewed as a single genetic subtype and patients are treated without regard to the EML4 fusion partner (Sabir et al. 2017; Schneider et al. 2023).

In addition to oncogenic drivers, tumors almost invariably have alterations in tumor suppressor genes that contribute to every aspect of tumorigenesis (Zehir et al. 2017; Martínez-Jiménez et al. 2020; Consortium et al.). While EML4-ALK-driven tumors tend to have lower mutational burdens than other subtypes of lung cancer, most contain alterations in genes that function as tumor suppressors in other contexts (Noh et al. 2017; Lee et al. 2019). However, beyond the association of *TP53* mutations with more aggressive disease, little is known about the impact of tumor suppressor genes in EML4-ALK-driven lung adenocarcinoma (Kron et al. 2018; Christopoulos et al. 2018; Zhang et al. 2021). Moreover, the genomic landscape of EML4-ALK-driven lung cancer is relatively poorly characterized due to its rarity and many existing cancer genomic datasets do not differentiate between EML4-ALK variants.

Our understanding of EML4-ALK in lung adenocarcinoma has been facilitated by pre-clinical studies in cell lines and mouse models (Soda et al. 2007; Gainor et al. 2016; Recondo et al. 2020; Chen et al. 2022; Shiba-Ishii et al. 2022; Diaz-Jimenez et al. 2024). In particular, autochthonous mouse models recapitulate key features of human tumors and have contributed to our understanding of the genetic drivers of human cancer. Overexpression of human *EML4-ALK* V1 and CRISPR/Cas9-mediated generation of the *Eml4-Alk* V1 rearrangement in mouse lung epithelial cells have been used to investigate V1-driven lung tumorigenesis *in vivo* (Soda et al. 2008; Blasco et al. 2014; Maddalo et al. 2014; Pyo et al. 2017). However, no other EML4-ALK variants have been modeled *in vivo*, and existing models have not been used to illuminate the impact of coincident genomic alterations. Leveraging high-throughput *in vivo* functional genomic methods to map interactions between putative tumor suppressors and EML4-ALK variants could thus elucidate potential differences between EML4-ALK variants and provide unique insights into pathways that influence EML4-ALK-driven tumor growth.

Here, we integrate tumor barcoding with novel autochthonous mouse models of EML4-ALK V1 and V3-driven lung cancer. We identify and validate diverse genetic interactions where alterations in tumor suppressor genes strongly and specifically influence tumorigenesis driven by either EML4-ALK V1 or V3. These results, together with molecular studies of V1- and V3-driven tumors and analysis of a large, novel cohort of lung cancer patients with EML4-ALK fusions, suggest that oncogenic fusion partners dramatically impact the effect of coincident genomic alterations on tumor development.

## RESULTS

### Lentiviral CRISPR/Cas9-based modeling of EML4-ALK V1- and V3-driven lung tumorigenesis combined with coincident gene inactivation

Lentiviral vectors stably integrate into the genomes of transduced cells, enabling barcoding-based clonal analyses in autochthonous tumor models (Rogers et al. 2017; Cai et al. 2021; Langille et al. 2022; Blair et al. 2023; Dervovic et al. 2023; Tang et al. 2023). To determine whether lentiviral vectors can induce EML4-ALK V1- and V3-driven lung tumors through inversion of the endogenous genomic region, we generated vectors encoding Cre recombinase and sgRNAs targeting introns of *Eml4* (intron 14 for V1 and intron 6 for V3) and intron 19 of *Alk* (Lenti-sgV1/Cre and Lenti-sgV3/Cre) (**Figure 1a** and **Supplementary Figure S1a,** (Maddalo et al. 2014; Diaz-Jimenez et al. 2024)). Transduction of lung epithelial cells in *R26^LSL-Cas9-EGFP^* mice (hereafter, *Cas9-EGFP*, (Platt et al. 2014)) with these vectors induced expression of Cas9 in addition to sg*Eml4* and sg*Alk,* resulting in double-stranded DNA breaks at the targeted loci, chromosomal inversion, and expression of the *Eml4-Alk* fusion oncogenes (**Figure 1a** and **Supplementary Figure S1a,b**).

**Figure 1.**
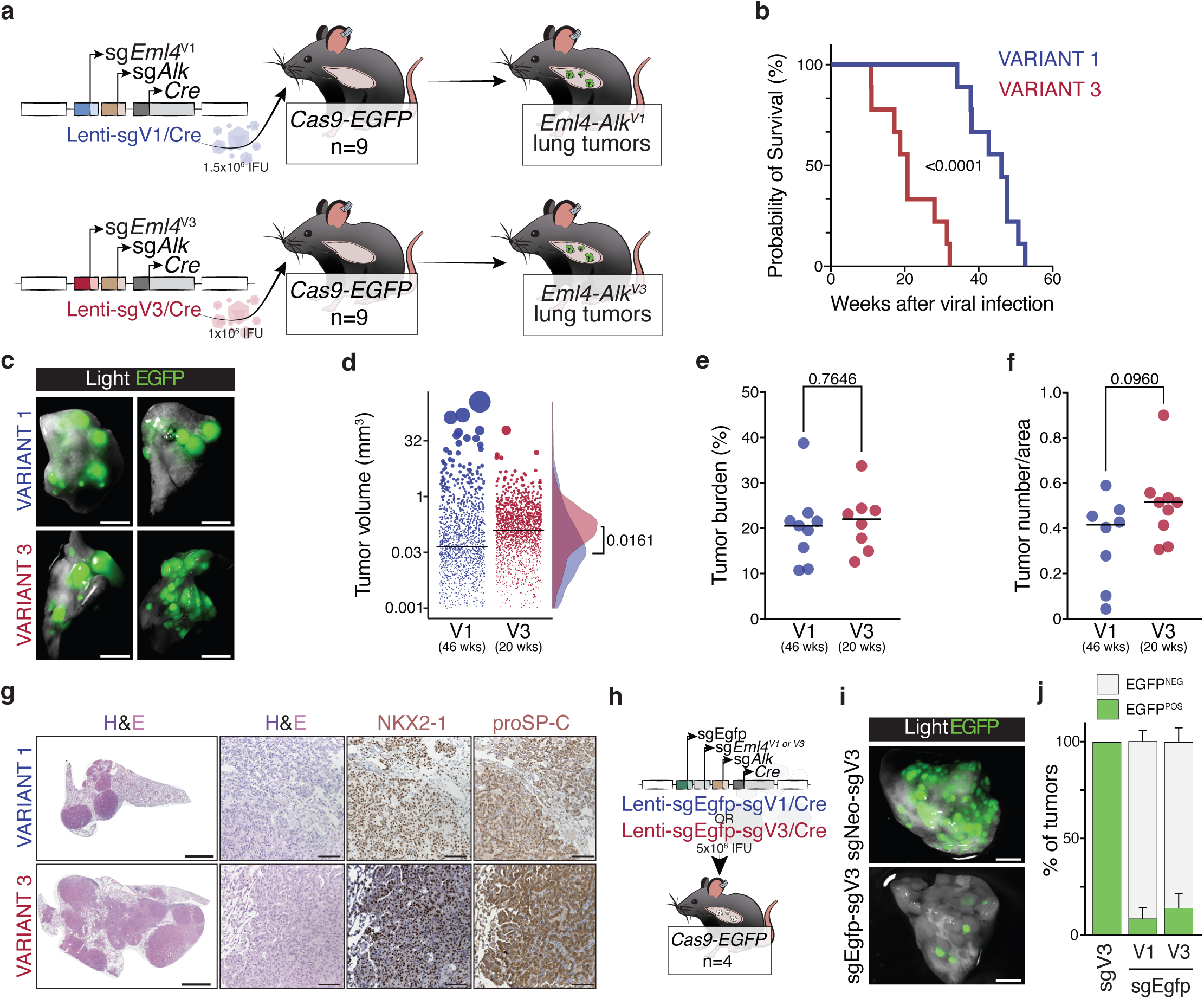
Lentiviral CRISPR/Cas9-mediated initiation of *Eml4-Alk* V1 and V3 lung tumors and efficient coincident gene inactivation. **a.** Schematic of CRISPR/Cas9-mediated initiation of *Eml4-Alk* variant 1 (V1)- and variant 3 (V3)-driven lung tumors in *R26^LSL-Cas9-EGFP/+^* (*Cas9-EGFP*) mice by intratracheal injection of Lenti-sgV1/Cre and Lenti-sgV3/Cre lentiviral vectors. Viral titer per mouse, mouse number, and genotype are indicated. **b.** The overall survival of *Cas9-EGFP* mice with V1 or V3 tumors is presented, with a median survival of 46 weeks for V1 (n=9) and 20.57 weeks for V3 (n=9). A Log-rank (Mantel-Cox) test shows a p-value of <0.0001. **c.** Light and fluorescent images show EGFP-positive tumors in the lungs of mice injected with Lenti-sgV1/Cre and Lenti-sgV3/Cre vectors. Scale bars are 2 mm. **d.** Individual *Eml4-Alk* V1 and V3 tumor volumes. The marginal density plots show the distribution of V1 and V3 tumor volumes along the Y-axis. An unpaired t-test was performed, and the nominal P-value is displayed. **e.** Tumor burden (Tumor area/Total area, %) from end-point mice with V1 or V3 tumors. Each dot represents a mouse. Unpaired t-test, nominal P-value is shown. **f.** Tumor numbers in mice with V1 or V3 tumors. Each dot represents a mouse. Unpaired t-test, nominal P-value is shown. **g.** Representative H&E staining and immunohistochemistry for NKX2-1 and proSP-C of lung sections from mice with V1 and V3 lung tumors. Higher magnification of V1 and V3 tumors with high-grade adenocarcinoma features. Scale bars are 1 mm, 100 µm, and 10 µm. **h.** Schematic of *Cas9-EGFP* mice intratracheally injected with Lenti-sgEgfp-sgV1/Cre or Lenti-sgEgfp-sgV3/Cre vectors. Viral titer per mouse, mouse number, and genotype are indicated. **i.** Light and fluorescent images from EGFP-positive lung tumors in mice injected with Lenti-sgNeo-sgV3/Cre or Lenti-sgEgfp-sgV3/Cre vectors. Scale bars are 1 mm. **j.** Percentage of EGFP-positive and negative tumors in *Cas9-EGFP* mice injected wtih Lenti-sgEgfp-sgV1/Cre or Lenti-sgEgfp-sgV3/Cre vectors. Error bars represent standard deviation.

All *Cas9-EGFP* mice developed lung tumors after intratracheal viral delivery of Lenti-sgV1/Cre or Lenti-sgV3/Cre. However, mice transduced with Lenti-sgV3/Cre had significantly shorter survival (∼20 weeks) relative to mice transduced with Lenti-sgV1/Cre (∼46 weeks) (**Figure 1b**). While all mice had very high overall tumor burdens, V3-driven lung tumors were on average larger than those driven by V1 (**Figure 1c-f**). V1- and V3-driven lung tumors were adenomas and adenocarcinomas that expressed the alveolar epithelial lineage–defining transcription factor and classical lung adenocarcinoma marker NKX2-1/TTF1 as well as surfactant protein C (**Figure 1g**). Analysis of EGFP^positive^ neoplastic cells from micro-dissected V1 or V3 tumors confirmed the presence of the expected *Eml4-Alk* inversions (**Supplementary Figure S1c**). Delivery of Lenti-sgV1/Cre or Lenti-sgV3/Cre to mice with a different Cas9 allele (*R26^LSL-Tomato^;H11^LSL-Cas9^* mice, **Supplementary Figure S1b**, (Chiou et al. 2015)) also generated lung tumors and confirmed that V3 tumors are larger and more numerous than V1 tumors when analyzed at the same time point (**Supplementary Figure S1d-g**).

To enable coincident generation of the *Eml4-Alk* V1 or V3 fusion oncogene and inactivation of tumor suppressor genes of interest we modified Lenti-sgV1/Cre and Lenti-sgV3/Cre vectors to include a third sgRNA. We used U6 promoters from different species, modified tracrRNAs, and ordered the sgRNAs to minimize the impact of recombination and lentiviral template switching (**Methods** and **Supplementary Figure S1h,** (Murray et al. 2019)). To quantify the frequency of homozygous inactivation of the additionally targeted genes, we initiated tumors in *Cas9-EGFP* mice (that were homozygous for the *Rosa26^LSL-Cas9-EGFP^* allele) with triple sgRNA vectors containing the sgRNAs required to generate the *Eml4-Alk* V1 or V3 fusions as well as an sgRNA targeting *EGFP* (**Figure 1h**). More than 80% of the resulting Lenti-sgEgfp-sgV1/Cre and Lenti-sgEgfp-sgV3/Cre-initiated tumors completely lacked EGFP protein, suggesting efficient homozygous gene inactivation and *Eml4-Alk* rearrangement using this vector system (**Figure 1i,j** and **Supplementary Figure S1i**).

These results demonstrate our ability to engineer *Eml4-Alk* V1 and V3 fusion oncogenes and inactivate additional genes of interest using lentiviral vectors and somatic CRISPR/Cas9-mediated genome editing. Furthermore, these data suggest that V3 is a more potent oncogenic driver than V1.

### Quantification of the impact of tumor suppressor gene inactivation on V1 and V3 tumor growth

EML4-ALK*-*driven human lung tumors frequently have mutations in genes that suppress tumorigenesis in other cancer types, but have yet to be investigated in EML4-ALK-driven cancer (Kron et al. 2018; Christopoulos et al. 2018), **Supplementary Figure S2a** and **Supplementary Table 1**). To evaluate the role of tumor suppressor genes in EML4-ALK V1 and V3-driven lung cancer, we used a multiplexed approach that combined CRISPR/Cas9-mediated somatic genome editing and tumor barcoding coupled with high-throughput barcode sequencing to quantify the impact of 6 canonical lung tumor suppressor genes (Tuba-seq; **Figure 2a,b** (Cai et al. 2021; Blair et al. 2023). We generated two pools of barcoded lentiviral vectors encoding Cre, the V1 or V3 fusion-inducing *Eml4*- and *Alk*-targeting sgRNAs, and a variable third sgRNA targeting genes of interest (**Supplementary Figure S2b**). These pools included two to three sgRNAs targeting each gene of interest, five sg*Inert* control vectors, and an sgRNA targeting an essential gene (*Pcna*) (Lenti-sgTSG^19^-sgV1-BC/Cre and Lenti-sgTSG^19^-sgV3-BC/Cre, **Figure 2a**). Each vector contained a sgRNA identifier (sgID) and a random barcode (BC) sequence that uniquely labels each clonal tumor. We delivered Lenti-sgTSG^19^-sgV1-BC/Cre or Lenti-sgTSG^19^-sgV3-BC/Cre to the lungs of independent cohorts of *Cas9-EGFP* mice and Cas9-negative *Kras^LSL-G12D/+^* mice (**Figure 2b** and **Supplementary Figure S2c**). *Kras^LSL-G12D/+^* mice enable precise quantification of the proportion of each vector in each lentiviral pool, which is required to accurately calculate the effect of each sgRNA on tumorigenesis in the *Cas9-EGFP* mice (**Supplementary Figure S2c,d** and **Methods**). *Cas9-EGFP* mice were analyzed when they began to show signs of high lung tumor burden (∼14 and ∼8 weeks for mice with V1 and V3-driven tumors, respectively; **Figure 2b** and **Supplementary Figure S2e**). Despite being analyzed ∼6 weeks earlier after tumor initiation, mice with V3-driven tumors had higher lung weights than mice with V1-driven tumors, consistent with V3 being more oncogenic than V1 **(Figure 2c** and **Supplementary Figure S2f)**.

**Figure 2.**
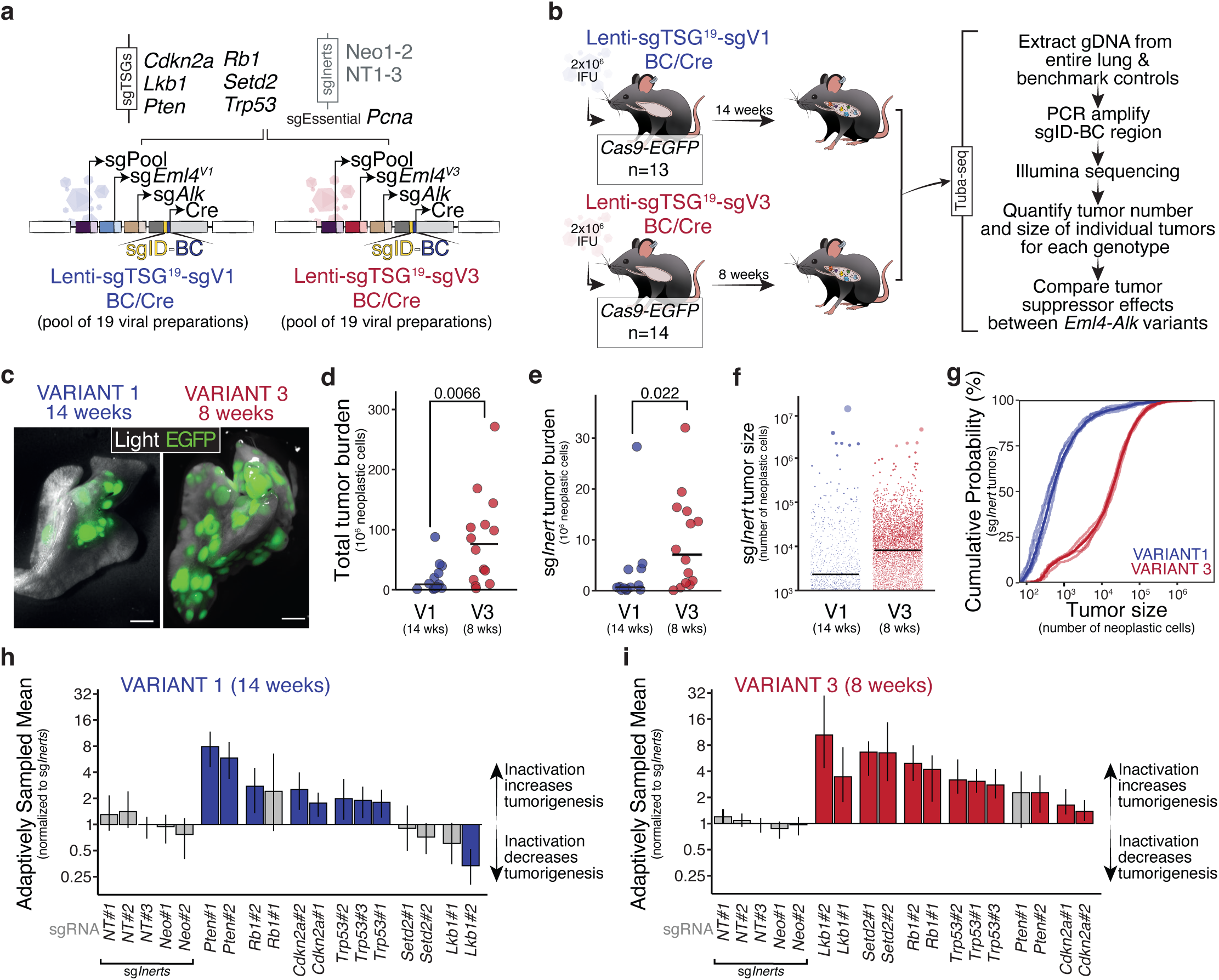
**EML4-ALK V1 and V3 differ in oncogenicity and are differentially impacted by tumor suppressor gene inactivation.** **a.** Lenti-sgTSG^19^-sgV1/Cre and Lenti-sgTSG^19^-sgV3/Cre. Pools include 2-3 barcoded vectors targeting 6 putative tumor suppressor genes of interest, 5 vectors encoding inert sgRNAs, and a vector with a sgRNA targeting an essential gene (*Pcna*). **b.** Initiation of lung tumors in *R26^LSL-Cas9-EGFP/+^*and *R26^LSL-Cas9-EGFP/LSL-Cas9-EGFP^ (Cas9-EGFP)* mice with Lenti-sgTSG^19^-sgV1/Cre (“V1 cohort”) and Lenti-sgTSG^19^-sgV3/Cre (“V3 cohort”). Lentiviral titer, median time-point of analysis, and mouse number are indicated. **c.** Fluorescent images of the lungs of mice from the V1 and V3 cohorts. EGFP fluorescence and bright-field images were merged. Scale bars: 2.5 mm. **d,e.** Total number of neoplastic cells summed across all tumors (**e**) and across all sgInert tumors in the V1 and V3 cohorts. Each dot represents a mouse and bars indicate median values within each genotype. The P-value was calculated using a two-sided Wilcoxon rank sum test. **f.** Visualization of *sgInert* tumor sizes in the V1 and V3 cohorts. Each dot represents a tumor and has an area proportional to its size. All tumors >1000 cells in size are plotted. Bars indicate the median tumor size within each cohort. **g.** Cumulative distribution functions of tumor size for adaptively sampled *sgInert* tumors in the V1 and V3 cohorts. Each translucent line represents an sgRNA; the solid lines are the cumulative density across sgRNAs. Note that the plot is truncated at 10^6^ cells to improve resolution at smaller tumor sizes. **h,i.** Adaptively sampled mean size (ASM) of each tumor genotype normalized to the ASM of *sgInert* tumors in the V1 (**h**) and V3 cohorts (**i**). ASM is a summary metric of tumor fitness that integrates the impact of inactivating each gene on tumor size and number. Each bar is a sgRNA. sgRNAs are grouped by gene. Baseline tumor growth (Y=1): no impact relative to *sgInert*. sgRNAs that significantly increase or decrease ASM (two-sided FDR-adjusted P-value) are in color. P-values and confidence intervals were calculated using nested bootstrap resampling.

To quantify the number and size of tumors with each sgRNA, we PCR-amplified and high-throughput sequenced the sgID-BC region of the integrated lentiviral vector from DNA extracted from the bulk tumor-bearing lungs. Mice with V3-driven tumors had greater tumor burden (total neoplastic cell number) and higher tumor number (number of barcoded clonal expansions with >1000 neoplastic cells) than mice with V1-driven tumors (**Figure 2d**, and **Supplementary Figure S2g,h**). Tumors initiated by sg*Inert* vectors (“sgInert tumors”) lack coincident gene inactivation and are driven solely by EML4-ALK V1 or V3. To quantify the oncogenicity of V1 and V3 independently of the effects of tumor suppressor gene inactivation, we compared the initiation and growth of sg*Inert* V1- and V3-driven tumors. sg*Inert* tumor burden and number were higher and median tumor size was larger in mice with V3-driven tumors relative to mice with V1-driven tumors (**Figure 2e,f** and **Supplementary Figure S2i**). To compare equivalent portions of the V1 and V3 tumor size distribution we developed a novel method which we term “adaptive sampling” in which the same number of tumors per unit of virus delivered are analyzed for each tumor genotype (**Supplementary Figure S2d** and **Methods**). This approach takes into account the number of mice, the lentiviral titer, and the relative proportion of each sgRNA vector in each pool (**Supplementary Figure S2c,d,** and **Methods**). This analysis confirmed that V3 tumors are substantially larger than V1 tumors, despite the longer duration of V1-driven tumor growth (**Figure 2g**).

We next analyzed the impact of tumor suppressor gene inactivation on V1- and V3-driven tumorigenesis. Using our adaptive sampling approach, we calculated the log-normal mean size (ASM) and the 95^th^ percentile tumor size (AS95) for each sgRNA in the V1 and V3 pools (**Figure 2h,i**, **Supplementary Figure S2j, S3a,b,** and **Methods**). The tumor-suppressive effects were generally consistent across sgRNAs targeting the same gene (consistent with on-target effects); therefore, we summarized the ASM and AS95 per gene by weighting each sgRNA in proportion to the number of adaptively sampled tumors (**Supplementary Figure S3c-f** and **Supplementary Table 2**). Inactivation of the essential gene *Pcna* reduced the ASM and AS95 by > 15-fold in both V1- and V3-driven tumors, confirming efficient coincident gene inactivation in both genetic contexts (**Supplementary Figure S3g,h**). As expected, sgRNAs had negligible effects in Cas9-negative *Kras^LSL-G12D/+^* mice, confirming that effects in the *Cas9-EGFP* mice are due to CRISPR/Cas9-mediated gene targeting (**Supplementary Figure S3i**,**j**). Inactivation of *Trp53* or *Cdkn2a*, which are frequently mutated or deleted in human EML4-ALK-driven lung tumors ((Kron et al. 2018; Christopoulos et al. 2018; Zhang et al. 2021), **Supplementary Table 1**), increased both V1-and V3-driven lung tumorigenesis (**Figure 2h,i**, **Supplementary Figure S3a-f**). The impact of inactivating several other tumor suppressor genes differed substantially between V1- and V3- driven tumors. Interestingly, the inactivation of *Pten* had a significantly larger effect on the growth of V1-compared to V3-driven tumors, while the inactivation of *Lkb1* or *Setd2* markedly increased V3-driven tumorigenesis but did not affect or even reduced V1-driven tumorigenesis (**Figure 2h,i**; **Supplementary Figure S3k)**.

### EML4-ALK V1 and V3-driven lung tumor growth is constrained by diverse tumor suppressor genes

To determine whether there are widespread differences in tumor suppressor function between V1- and V3-driven tumors, we next interrogated the effects of a broader panel of tumor suppressor genes. We prioritized 29 genes based on their mutation frequency in *EML4-ALK* lung cancer, their function in other lung cancer models, and the diversity of their functions and pathways (**Supplementary Figure S2a**, **Supplementary Table 1** and **Methods**, (Cai et al. 2021; Foggetti et al. 2021)). We generated additional Lenti-sgV1/Cre and Lenti-sgV3/Cre vectors such that we had two to four vectors with sgRNAs targeting each gene as well as six sg*Inert* vectors. Each of these 150 vectors was barcoded with a unique sgID-BC, followed by lentiviral production before pooling to generate Lenti-sgTSG^75^-sgV1-BC/Cre and Lenti-sgTSG^75^-sgV3-BC/Cre pools (**Figure 3a, Supplementary Figure S4a** and **Supplementary Table 3**).

**Figure 3.**
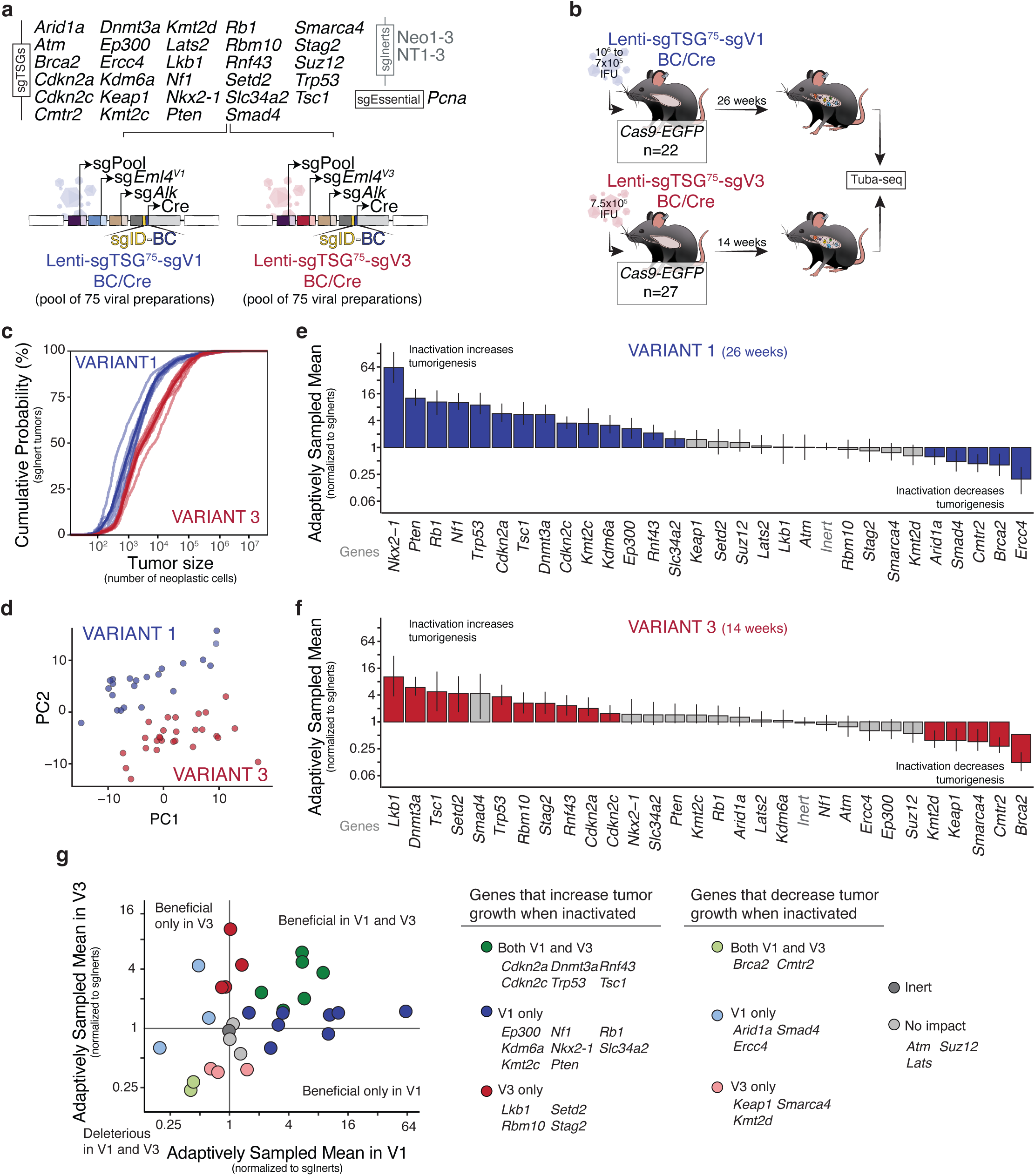
EML4-ALK V1 and V3-driven lung tumor growth is differentially constrained by diverse tumor suppressor genes. **a.** Lenti-sgTSG^75^-sgV1/Cre and Lenti-sgTSG^75^-sgV3/Cre pools include 2-4 barcoded vectors targeting each of the 29 putative tumor suppressor genes of interest, 6 vectors encoding inert sgRNAs, and a vector with a sgRNA targeting an essential gene (*Pcna*). **b.** Initiation of lung tumors in *R26^LSL-Cas9-EGFP/LSL-Cas9-EGFP^ (Cas9-EGFP)* mice with Lenti-sgTSG^75^-sgV1/Cre (”V1 cohort”) and Lenti-sgTSG^75^-sgV3/Cre (”V3 cohort”). Lentiviral titer, median time point of analysis, and mouse number are indicated. **c.** Cumulative distribution functions of tumor size for adaptively sampled sg*Inert* tumors in the V1 and V3 cohorts. Each translucent line represents one of the inert sgRNAs; the solid lines are the cumulative density across all inert sgRNAs. **d.** Principal components analysis of tumor suppressive effects performed within each mouse reveals consistent separation of mice with Lenti-sgTSG^75^-sgV1/Cre and Lenti-sgTSG^75^-sgV3/Cre initiate tumors. Each dot is a mouse. **e-f.** Adaptively sampled mean size (ASM) of each tumor genotype normalized to the ASM of sg*Inert* tumors in the V1 (**e**) and V3 cohorts (**f**). ASM is a summary metric of tumor fitness that integrates the impact of inactivating each gene on tumor size and number. Baseline tumor growth (Y=1): no significant impact. Genes that significantly increase or decrease ASM (two-sided FDR-adjusted p-value) are in color. P-values and confidence intervals were calculated using nested bootstrap resampling. **g.** Summary of the impact of all genes assayed in the Lenti-sgTSG^75^ pools on V1- and V3-driven tumorigenesis. Left, comparison of ASM for all tumor genotypes in the V1 and V3 cohorts. The impact of *Pcna* inactivation is not shown to improve visualization of smaller magnitudes of effect. Genes are colored as indicated on the right. Right, categorization of genes as tumor suppressor genes (TSG) or tumor-promoting genes in V1- and/or V3-driven tumorigenesis. TSG: ASM > 1 and two-sided FDR-adjusted P-value < 0.05. Tumor-promoting gene: ASM < 1 and two-sided FDR-adjusted P-value < 0.05.

We initiated tumors with Lenti-sgTSG^75^-sgV1-BC/Cre and Lenti-sgTSG^75^-sgV3-BC/Cre in separate cohorts of *Cas9-EGFP* mice as well as in cohorts of Cas9-negative control mice (**Figure 3b** and **Supplementary Figure S4b**). Lungs were collected for Tuba-seq when mice showed signs of high tumor burden, resulting in similar lung weights, tumor burdens, and tumor numbers in *Cas9-EGFP* mice transduced with the Lenti-sgTSG^75^-sgV1-BC/Cre and Lenti-sgTSG^75^-sgV3-BC/Cre pools (∼26 and ∼14 weeks for mice with V1 and V3-driven tumors, respectively; **Supplementary Figure S4c-j**). Comparison of the size distributions of adaptively sampled sg*Inert* tumors confirmed that V3-driven tumors were larger than V1-driven tumors (**Figure 3c**). Principal component analysis of the relative size of tumors at the 95^th^ percentile of the tumor size distribution for each sgRNA distinguished mice with V1-driven tumors from those with V3-driven tumors, suggesting that there are consistent differences in the effects of inactivating these genes in V1- and V3-driven tumors (**Figure 3d**).

To quantify the impact of each tumor suppressor gene on V1- and V3-driven tumorigenesis, we calculated the adaptively sampled log-normal mean size (ASM) and the adaptively sampled 95^th^ percentile tumor size (AS95) for each sgRNA and gene (**Figure 3e**,**f**, **Supplementary Figure S5** and **Supplementary Table 4**). V1- and V3-driven lung tumorigenesis *in vivo* were extensively impacted by coincident tumor suppressor gene inactivation. For each EML4-ALK variant, some genotypes dramatically increased tumorigenesis, whereas others decreased tumorigenesis (**Figure 3e-g**). Roughly equal numbers of genes were tumor suppressors in V1 and V3 tumors, but the magnitude of tumor-suppressive effects was larger in V1 tumors, consistent with diminishing returns epistasis, a phenomenon in which the fitness advantage conferred by a genetic alteration decreases on a more fit genetic background (**Figure 3e-g**, (Wei and Zhang 2019)). The effects of sgRNAs targeting the same gene were largely similar (**Supplementary Figure S5a,c**). Consistent with our initial screen, *Trp53* and *Cdkn2a* suppressed both V1 and V3-driven lung tumorigenesis (**Figure 3e-g**). Interestingly, the inactivation of some genes that are generally considered tumor suppressors reduced tumor growth. For example, inactivation of *Smad4* reduced V1-driven growth, and inactivation of *Cmtr2* reduced V1- and V3- driven growth (**Figure 3e-g**). Several genes not assayed in our initial screen were identified as critical tumor suppressors, with inactivation of *Nkx2-1* or *Nf1* dramatically increasing V1-driven tumorigenesis and inactivation of *Dnmt3a* or *Tsc1* increasing both V1- and V3-driven tumorigenesis (**Figure 3e-g**).

To quantify differences in tumor suppressor function between V1- and V3-driven tumors, we compared the ASMs for each genotype in V1 and V3. The effects of inactivating the majority of genes differed between these oncogenic contexts, suggesting that the landscapes of tumor suppression in V1- and V3-driven lung tumorigenesis are strikingly different (**Supplementary Figure S4k**).

### Validation of variant-specific effects of tumor suppressor gene inactivation

To validate the differential effects of inactivating *Pten*, *Rb1*, and *Setd2* on V1- and V3- driven tumors outside of the pooled setting, we initiated tumors in eight separate cohorts of *Cas9-EGFP* mice with individual sg*Pten*, sg*Rb1*, sg*Setd2* and sg*Neo* expressing Lenti-sgV1/Cre and Lenti-sgV3/Cre vectors (**Figure 4a**). Consistent with its greater effect in the Lenti-sgTSG^75^-sgV1-BC/Cre pool, *Rb1* inactivation significantly decreased the survival of mice with V1-but not V3-driven tumors. Likewise, while *Setd2* inactivation decreased the survival of mice with either V1 or V3-driven tumors, the effect was larger and more significant for V3-driven tumors, consistent with the Tuba-seq analysis. *Pten* inactivation reduced survival of mice with V1- and V3-driven tumors, although this did not reach significance (**Figure 4b**). Quantification of surface tumor sizes from all samples and from a subset of samples that were collected at similar time points confirmed variant-specific interactions, with *Pten* or *Rb1* inactivation significantly increasing V1-driven tumor size and *Setd2* inactivation significantly increasing V3-driven tumor size (**Figure 4c** and **Supplementary Figure S6a,b**). Through immunohistochemistry for the SETD2-deoendent histone mark H3K36me3, we confirmed *Setd2* inactivation in the majority of V1 and V3 tumors initiated with Lenti-sg*Setd2*/Cre (**Supplementary Figure S6c,d**). Tumors lacking H3K36me3 (*Setd2*-deficient tumors) were significantly larger than tumors with H3K36me3 (*Setd2*-proficient tumors) within individual mice (**Supplementary Figure S6e**). This effect was more consistent and dramatic in V3-driven tumors, further suggesting that SETD2 is a stronger tumor suppressor in V3-driven tumors (**Supplementary Figure S6e**).

**Figure 4.**
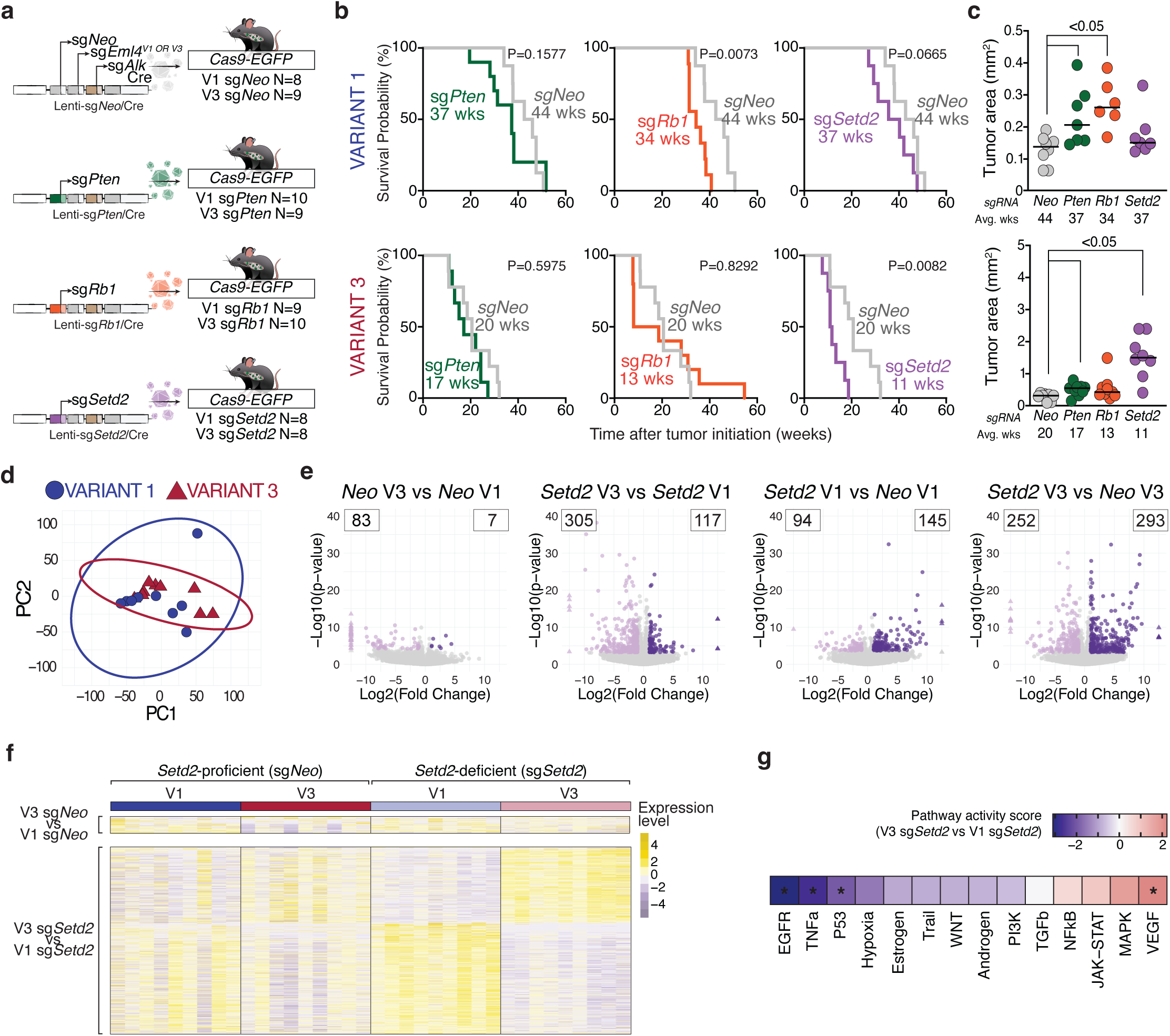
Validation experiments and molecular analyses of EML4-ALK V1 and V3-driven tumors confirm variant-specific effects of tumor suppressor gene inactivation. **a.**Schematic of tumor initiation in *R^26LSL-Cas9-EGFP/+^* (*Cas9-EGFP)* mice with individual Lenti-sgTSG-sgV1/Cre and Lenti-sgTSG-sgV3/Cre vectors. Titers were 1.5x10^6^ IFU/mouse for all Lenti-sgTSG-sgV1/Cre vectors and 1x10^6^ IFU/mouse for all Lenti-sgTSG-sgV3/Cre vectors. **b.** Overall survival of *Cas9-EGFP* mice transduced with Lenti-sgTSG-sgV1/Cre and Lenti-sgTSG-sgV3/Cre vectors. Grey line indicates sgNeo-V1 or sgNeo-V3 mice; colored lines indicate sgTSG-V1 or sgTSG-V3 mice. Median survival for each group is indicated. Log-rank (Mantel-Cox) was performed and nominal P-values are indicated. **c.** Surface tumor area (mm^2^) from indicated tumor genotypes. Each point represents the median tumor size in one mouse. Horizontal lines indicate the mean within each group. Nominal P-values for significant comparisons versus Neo are indicated. **d.** Principal components analysis using the 6,000 most variable genes from sgNeo-V1 and sgNeo-V3 mice. Each point represents the gene expression profile of cancer cells derived from a single tumor. Points are colored according to their *Eml4-Alk* variant. **e.** Volcano plots showing changes in gene expression for indicated contrasts.The number of up- or downregulated genes is indicated. Triangles indicate gene with LFC >|10|. **f.** Heatmaps showing Z-scored normalized counts for differentially expressed genes from indicated contrasts across all *Setd2*-deficient and *Setd2*-proficient V1- and V3-driven tumors. **g.** Inferred pathway activity scores comparing *Setd2*-deficient V3 (V3 sg*Setd2*) to *Setd2*-deficient V1 (V1 sg*Setd2*) tumors. The inferred activity is based on the t-values of the differentially expressed genes for each comparison. Positive scores (red) indicate that a pathway is more active in V3 sg*Setd2* tumors; negative scores (blue) indicate that a pathway is more active in V1 sg*Setd2*. Stars denote that a pathway is differentially activated (p<0.05, PROGENy multivariate linear model).

*TP53* is frequently mutated in human lung adenocarcinomas regardless of their EML4-ALK variant and our Tuba-seq data suggested that P53 suppresses both V1- and V3-driven lung tumors. We initiated *Trp53*-deficient V1- and V3-driven tumors in *Trp53^flox/flox^;Cas9-EGFP* mice with Lenti-sgV1/Cre and Lenti-sgV3/Cre vectors (**Supplementary Figure S6f**). As anticipated, inactivation of *Trp53* enhanced the growth of both V1- and V3-driven tumors (**Supplementary Figure S6g-i**). Collectively, these data validate that there are epistatic interactions between specific tumor suppressor genes and EML4-ALK fusion variants.

### EML4-ALK V1- and V3-driven tumors have distinct transcriptional responses

Our *in vivo* functional data demonstrate that the EML4 fusion partner determines the extent to which many tumor suppressor genes impact tumorigenesis. To investigate the baseline molecular difference between EML4-ALK V1 and V3 tumors, as well as their molecular responses to the same genetic perturbation, we performed bulk RNA-sequencing on sorted EGFP^positive^ neoplastic cells from V1 and V3-driven tumors that were either *Setd2*-proficient (sg*Neo*) or *Setd2*-deficient (sg*Setd2*; **Figure 4a** and **Supplementary** Figure 7a-c). We chose to use *Setd2* inactivation as our perturbation not to study *Setd2* itself, but because it is frequently inactivated in EML4-ALK-driven tumors and has a variant-specific impact on tumor fitness (**Figures 2,3,4c** and **Supplementary Figure S6a-e**). Neoplastic cells from *Setd2*-proficient V1 and V3 tumors had similar gene expression profiles, with only a modest number of significantly differentially up- and down-regulated genes (**Figure 4d-f**). While *Setd2* deficiency generated gene- and pathway-level changes within both V1- and V3-driven tumors, V3-driven tumors had more widespread changes in expression (**Figure 4e** and **Supplementary Figure S7d,e**). Interestingly, Setd2 inactivation differentiated the gene expression profiles of V1- and V3-driven tumors, with neoplastic cells from *Setd2*-deficient V1 and V3 tumors having dramatic and consistent differences in gene expression (**Figure 4e-g**). Thus, lung tumors driven by different EML4-ALK variants have distinct transcriptional responses to a functional perturbation (in this case inactivation of *Setd2*), consistent with the differential impact of *Setd2* inactivation on tumor growth and more generally with the overall change in the landscape of tumor suppression.

### EML4-ALK V1 and V3 resemble distinct oncogenes in their tumor suppressive effects

The extent of difference between the effects of tumor suppressor inactivation in V1- and V3-driven tumors is striking given the presumed similarities between the two fusion proteins as oncogenic drivers. To further investigate this finding and facilitate comparison with existing datasets on tumor suppressor-oncogene epistasis, we calculated the Spearman’s rank correlation coefficient (ρ) for all genes in the Lenti-sgTSG^75^-sgV1-BC/Cre and Lenti-sgTSG^75^-sgV3-BC/Cre pools. The impacts of tumor suppressor gene inactivation were only weakly correlated between V1 and V3 (ρ=0.36, **Figure 5a, Supplementary Figure S8a,b** and **Methods**). Bootstrap resampling analyses confirmed that this was not driven by variability in our estimation of tumor suppressor effects in the V1 and V3 datasets (**Figure 5a**, **Supplementary Figure S8a,b** and **Methods**). Repeating our analyses using different numbers of adaptively sampled tumors and ranking genes based on AS95 rather than the ASM consistently showed that the impacts of tumor suppressor gene inactivation in V1- and V3-driven tumors were either weakly correlated or entirely uncorrelated (**Supplementary Figure S8c,d**).

**Figure 5.**
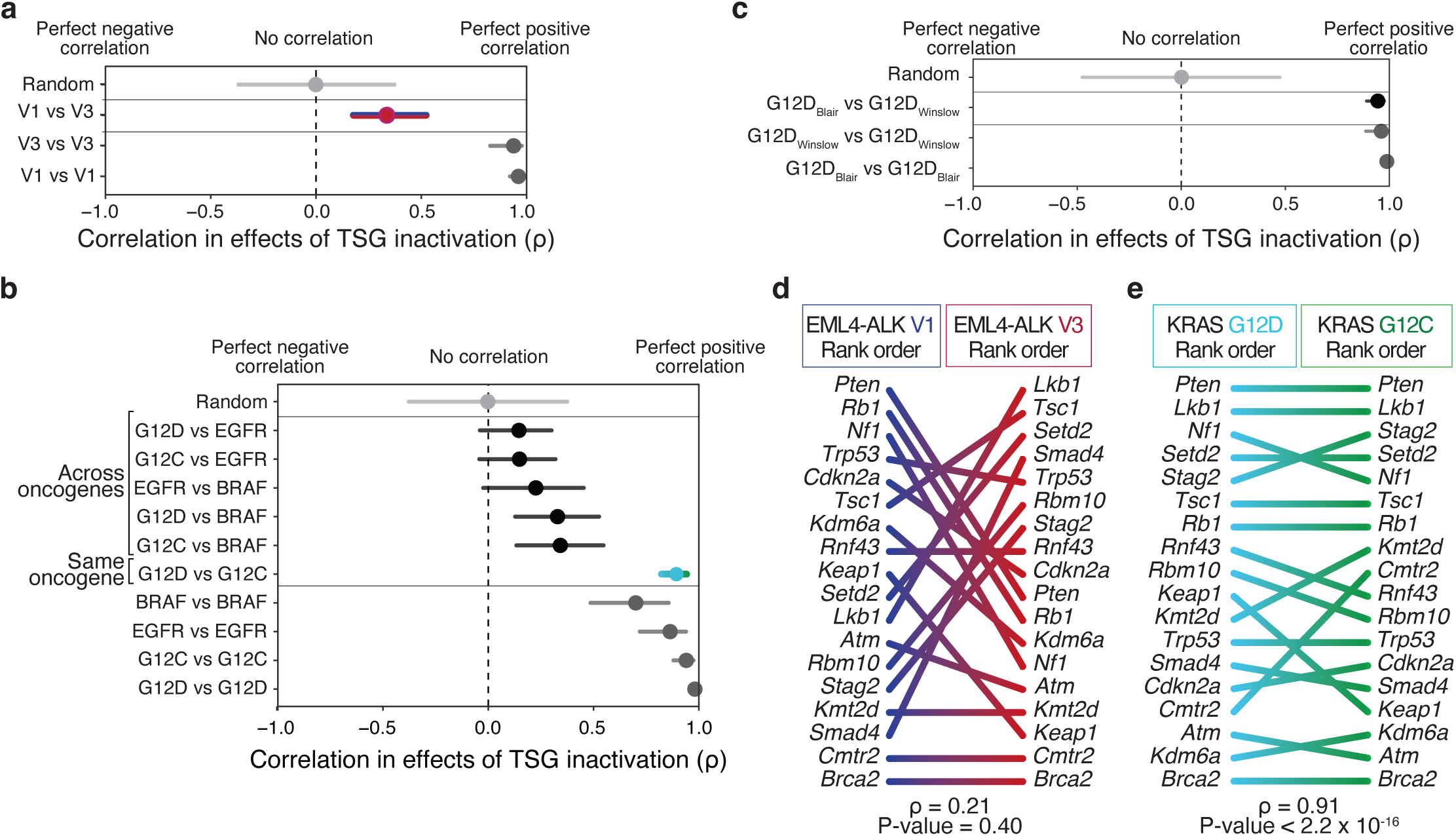
EML4-ALK V1 and V3 resemble distinct oncogenes with respect to their tumor suppressive landscapes. **a.** Correlation in the rank order of tumor suppressive effects in *Cas9-EGFP* mice transduced with the Lenti-sgTSG^75^-sgV1/Cre (V1) and Lenti-sgTSG^75^-sgV3/Cre (V3). Genes were ranked by fold change in ASM relative to sg*Inert* vectors. For panels **a**-**c**: Confidence intervals were generated through a nested bootstrap resampling procedure where tumor suppressive effects and ranks were repeatedly recalculated to quantify uncertainty in ρ. Points are median values across 10,000 bootstrap resamplings. “Random” reflects the simulated distribution of ρ observed when there is no relationship between two groups given the number of items being ranked. **b.** Correlation in the rank order of tumor suppressive effects between KRAS G12D, KRAS G12C, BRAF, and EGFR mice from Blair *et al*. Genes were ranked by fold change in 95^th^ percentile tumor size relative to sg*Inert*. **c.** Correlation in the rank order of tumor suppressive effects for KRAS G12D mice from Blair et al (”G12D_Blair_”) and an independent KRAS G12D dataset (”G12D_Winslow_”). Only the subset of genes assayed in both datasets were included (N=18). For both datasets, genes were ranked by fold change in 95 percentile tumor size relative to sg*Inert* vectors. **d,e.** Comparisons of the rank order of tumor suppressor effects between EML4-ALK V1 and V3 lung tumors (**d**) and between KRAS G12D and KRAS G12C lung tumors (data from Blair *et al*., **e**). Genes assayed in both the Lenti-[sgTSG-sgV1/V3]^Pool^BC/Cre pools and in Blair *et al*. are shown.

To contextualize these findings, we compared the correlation in tumor-suppressive effects between EML4-ALK V1 and V3 with analogous data from mouse models of oncogenic KRAS^G12D^, KRAS^G12C^, EGFR^L858R,^ and BRAF^V600E^-driven lung cancer (**Figure 5b** and **Methods**, (Blair et al. 2023)). For all comparisons between distinct pairs of oncogenes (i.e., a *Kras* variant with *Braf*, *Braf* with *Egfr*, or *Egfr* with a *Kras* variant), the impacts of tumor suppressor gene inactivation were only weakly correlated. In contrast, effects were extremely well correlated between oncogenic KRAS^G12D^- and KRAS^G12C^-driven tumors. We repeated these analyses and our analysis of the V1 and V3 datasets using only the subset of genes included in both the Lenti-sgTSG^75^ viral pools and the published data and found similar results (**Supplementary** Figure 8e,f). Importantly, the rank order of tumor suppressors in the published KRAS^G12D^ data (Blair et al. 2023) correlated extremely well with an independent KRAS^G12D^ dataset that we generated, suggesting that this ranking approach is effective in comparing tumor-suppressive effects across independent datasets (**Figure 5c**, (Shuldiner et al. 2024)). Visual comparison of the ordering of tumor suppressor genes when ranked on their effects in EML4-ALK V1 and V3 highlights the extensive reordering between these contexts and demonstrates that the poor correlation between them is driven by differences in the effects of strong tumor suppressors (**Figure 5d**). In contrast, both the frequency and magnitude of rank changes are greatly reduced between KRAS^G12D^ and KRAS^G12C^, with the same set of genes functioning as strong tumor suppressors in both contexts (**Figure 5e**).

Collectively, these analyses show that the extent of tumor suppressor epistasis in the V1 and V3 genetic backgrounds is qualitatively similar to patterns of epistasis across oncogenes and greatly exceeds that which has been observed between variants of the same oncogene. Our data show that at the level of their responses to the impact of diverse tumor suppressor genes EML4-ALK V1 and V3 more closely resemble distinct oncogenes than variants of the same oncogene (**Figure 5d,e** and **Supplementary Figure S8g-k**).

### Variant-resolved analysis of the genomic landscape of human EML4-ALK*-*driven lung cancer

The genomic landscape of human lung cancer driven by different EML4-ALK variants is poorly characterized due to the scarcity of patient-derived samples and inconsistent annotation of fusion variants in some databases. To address these limitations, we evaluated the largest cohort to date of non-small cell lung cancers with EML4-ALK fusions for which uniform genomic profiling was performed by Foundation Medicine, Inc (n= ∼2,000). The frequencies of EML4-ALK variants were consistent with prior clinical datasets, with most patients having V1 (34.2%) or V3 (35.7%) rearrangements (**Figure 6a** and **Supplementary Figure S9a**). Sequencing of 296 cancer-related genes revealed numerous recurrent alterations but low overall mutation burden (**Supplementary Table 5**). Consistent with prior data, the most common alterations were inactivating mutations in *TP53* and deletions of *CDKN2A* and *CDKN2B* (**Figure 6b**, (Kron et al. 2018; Christopoulos et al. 2018; Zhang et al. 2021). All other genes, including many of those targeted in the Lenti-sgTSG^75^ viral pools, were mutated at much lower frequencies (**Figure 6b**, **Supplementary Figure S9b**, and **Supplementary Table 5**). More than 75% percent of samples had alterations in known tumor suppressor genes, and almost half had alterations beyond those in *TP53*, *CDKN2A* and *CDKN2B*, highlighting the potential importance of this class of alteration in EML4-ALK-driven lung cancer (**Supplementary Figure S9c**). Interestingly, we identified genes with recurrent alterations that have not previously been described in EML4-ALK-driven lung tumors (**Supplementary Figure S9d**). Notably, amplifications of *ARFRP1*, a gene encoding a member of the ARF family of GTPases, occurred in 51 samples (2.6%). *ARFRP1* amplifications have been described in breast and colorectal cancers (Consortium et al. 2017), but are very rare (<0.5%) in non-small cell lung cancer in general and have not been described in previous EML4-ALK patient datasets. The relatively high prevalence of these alterations suggests that they may have an underappreciated importance in EML4-ALK-driven lung cancer.

**Figure 6.**
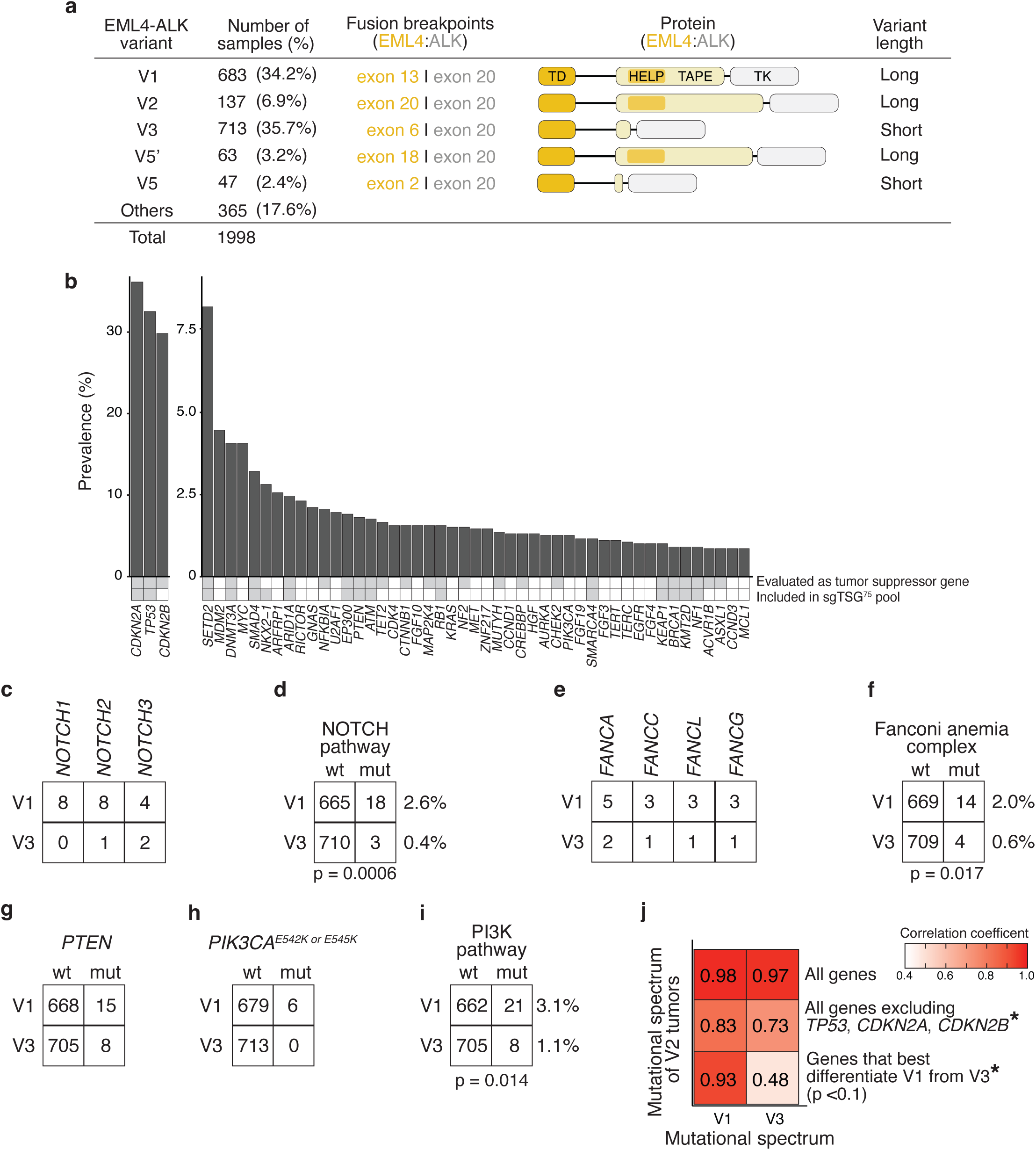
The genomic landscape of EML4-ALK-driven lung cancer. **a.**Number of samples per fusion variant in the EML4-ALK non-small cell lung cancer (NSCLC) patient cohort. Variants are annotated per short or long isoform. Schematics of the human loci (indicated relevant exons) and protein product are shown for each variant. EML4 and ALK relevant domains are indicated (trimerisation domain (TD); hydrophobic motif in EML proteins (HELP); tandem atypical propeller domain (TAPE); tyrosine kinase domain (TK)). **b.** Alteration frequencies across all samples in the EML4-ALK NSCLC patient cohort for the 50 most commonly altered genes. Annotations indicate genes evaluated as tumor suppressors in analysis of patient samples (top) and included in the sgTSG^75^ pool. **c-f**. Alterations in *NOTCH* genes (**c,d**) and genes encoding members of the Fanconi anemia complex (**d,f**) occur significantly more frequently in EML4-ALK V1-driven tumors relative to EML4-ALK V3-driven tumors. Two-sided Fisher’s Exact test p-values are reported. **g-i**. Alterations in *PTEN* and canonical oncogenic alterations of *PIK3CA* (*PIK3CAE^542K^* and *PIK3CAE^545K^*) occur significanly more frequency in EML4-ALK V1-driven tumors relative to EML4-ALK V3-driven tumors. Two-sided Fisher’s Exact test p-value is reported. **j.** The mutational spectrum of EML4-ALK V2-driven tumors more closely resembles that of EML4-ALK V1-driven tumors than EML4-ALK V3-driven tumors. Shading reflects Pearson correlation coefficient between alteration frequencies in EML4-ALK V2-driven tumors and alteration frequencies in EML4-ALK V1- and V3-driven tumors for the indicated gene sets. * indicates a signicant difference (two-sided Fisher’s z-test p<0.05) between correlation of V2-driven tumors with V1- and V3-driven tumors.

Given the widespread differences between the functional effects of tumor suppressor gene inactivation in V1- and V3-driven tumors (**Figures 3-5**), we hypothesized that the mutational spectrum of human tumors would vary depending on the EML4-ALK fusion variant. Tumor mutational burden was similar but slightly higher in V3-driven tumors relative to V1-driven tumors (**Supplementary Figure S9e**). Alteration frequencies were generally similar between V1- and V3-driven tumors and no genes had statistically significant differences in mutation frequency across variants after FDR correction (**Supplementary Tables 6, 7**). However, given the very low alteration frequencies of most genes, even with this dataset that includes > 10 times as many patients as described previously, we were underpowered to detect variant-level differences (**Supplementary Figure S9f**). To partially overcome this, we aggregated alteration counts across genes with related biological functions. This analysis identified several gene sets that are differentially mutated in tumors with each variant. Most significantly, *NOTCH* genes were > 7 times more likely to be mutated in V1-than V3-driven tumors (**Figure 6c,d**). Likewise, genes encoding members of the Fanconi anemia core complex were ∼3 times more likely to be mutated in V1-driven tumors (**Figure 6e,f**). Consistent with our functional in *vivo* data, mutations in *PTEN* occurred more frequently in V1-driven tumors, and canonical oncogenic alterations in *PIK3CA* were found exclusively in V1-driven tumors (**Figure 6g,h**). Collectively, these PI3K-activating alterations were significantly enriched and ∼3 times more frequent in V1-driven tumors (**Figure 6i**). Thus, while these data are limited in their ability to identify associations between individual genes and fusion variants, they independently support a model in which tumors driven by different EML4-ALK variants are biologically distinct and are differentially impacted by some tumor suppressor pathways.

Finally, our dataset includes patients with less common EML4-ALK variants, most notably the “long” EML4-ALK V2 (exon 20 of *EML4* fused with exon 20 of *ALK*; 6.9%, **Figure 6a** and **Supplementary Table 6**). Little is known about these variants due to their rarity; however, there may be similarities among variants based on the length of the EML4 partner protein (Sabir et al. 2017). Alteration frequencies in V2-driven tumors were highly correlated with alteration frequencies in both V1- and V3-driven tumors; however, this was largely driven by alterations in *TP53*, *CDKN2A*, and *CDKN2B*, which occur at high frequencies across all variants (**Figure 6j)**. Excluding these three genes revealed that the mutational spectrum of V2-driven tumors is significantly more correlated with the mutational spectrum of V1-driven tumors than V3-driven tumors (**Figure 6j**, p=0.005), consistent with the length of EML4 protein determining properties of the fusion variant. Interestingly, restricting our analysis to the subset of genes that best differentiate V1-driven tumors from V3-driven tumors (Fisher’s Exact test p<0.1, **Supplementary Table 7**) increased the correlation between V2- and V1-driven tumors while decreasing the correlation between V2 and V3, suggesting that there are consistent differences between the mutational spectra of tumors with “long” (*i.e.* V1 and V2) and “short” (V3) EML4-ALK variants (**Figure 6j**). In line with this, the patterns identified by comparing V1 and V3 were further reflected across tumors with V2, V5, and V5’ fusion variants. All *NOTCH* gene mutations in these less common variants occurred in samples with the “long” variant V2, and all mutations in *PTEN* occurred in samples with “long” variants V2 and V5’, further suggesting that differences between V1 and V3 map onto less commonly occurring “long” and “short” variants (**Supplementary Figure S9g**).

## DISCUSSION

Alterations in putative tumor suppressor genes are widespread in cancer, but the extent to which they constrain tumor initiation and progression in specific cancer types and how this is impacted by genomic context remains incompletely understood. Human cancer genomics, detailed cell biological experiments, and quantitative *in vivo* analyses have suggested that oncogenic context can strongly shape tumor suppressor function (Rogers et al. 2017; Cai et al. 2021; Langille et al. 2022; Blair et al. 2023; Dervovic et al. 2023; Tang et al. 2023). Here, we quantified the functional impact of inactivating a large and diverse panel of putative tumor suppressor genes on EML4-ALK V1 and V3-driven lung tumors, assessed molecular responses to tumor suppressor gene inactivation in both contexts, and investigated a novel genomic dataset from EML4-ALK-driven human lung tumors. We not only identified previously uncharacterized functional suppressors of EML4-ALK-driven lung tumors, but also uncover an unexpectedly complex fitness landscape where the identity of the EML4 fusion partner shapes the impact of important signaling pathways on tumor development.

EML4-ALK fusion oncogenes drive tumor growth through increased ALK kinase activity (Heuckmann et al. 2012; Childress et al. 2018; O’Regan et al. 2020). Consequently, one might have anticipated that the inactivation of diverse tumor suppressors would have similar biological consequences in V1- and V3-driven tumors. The prevalence and strength of variant-specific epistatic interactions that we observe challenge a simplistic model centered solely on ALK. The EML4-ALK V1 fusion includes the HELP domain and part of the TAPE domain of EML4 while the V3 fusion does not (**Figure 6a**). This difference has been suggested to impact the stability and functional properties of the EML4-ALK variants (Woo et al. 2016; Sabir et al. 2017; O’Regan et al. 2020). Given that V1 and V3 differ in the extent of EML4 domains, the tumor suppressor genes with similar effects in V1 and V3 could impact pathways predominantly downstream of ALK kinase activity, whereas those with different effects could operate predominantly downstream of the HELP or TAPE domains from EML4 (**Figure 6a**). Interestingly, V1, but not V3 has been shown to interact with RAS family members through the HELP domain (Hrustanovic et al. 2015). Consistent with these results, our findings show that inactivation of *Nf1*, a negative regulator of RAS, specifically increased the growth of V1 lung tumors. Our analysis of the genomic landscape of human EML4-ALK lung cancer identified similarities in mutational spectrum based on variant length, suggesting that the functional differences between V1 and V3 may extend to other EML4-ALK variants. Further characterization of additional EML4-ALK variants could assess this possibility and help to pinpoint the differences between V1 and V3 that drive differential effects of tumor suppressor inactivation.

Tumor suppressor genes regulate pathways and cellular processes that are critical to cancer development (Hanahan 2022). Our findings demonstrate that the consequences of perturbing these genes differ between V1- and V3-driven tumors at the level of both tumor fitness and gene expression, suggesting that there are fundamental differences in the biology of V1 and V3 tumors. These differences alone may impact responses to diverse therapies. Additionally, the inactivation of certain tumor suppressor genes could activate bypass pathways in a variant-dependent manner, further influencing therapy responses. These differences in the contributions of other signaling pathways to the growth of EML4-ALK V1 and V3-driven tumors may be meaningful for treatments that combine ALK inhibition with other therapies and for second-line therapies that are independent of ALK inhibition. Trials that do not incorporate data on the specific EML4-ALK variants and coincident tumor suppressor gene alterations could fail to uncover patient subsets that receive meaningful clinical benefit.

Lastly, our methodology has the capacity to generate lung tumors driven by specific EML4-ALK variants while simultaneously inactivating any panel of genes of interest, including additional tumor suppressor genes, biological pathways of interest, or candidate drug targets. These types of pre-clinical models also allow for the precise investigation of the response of specific tumor genotypes to therapies (Li et al. 2021; Foggetti et al. 2021), offering a potentially valuable system for the design and optimization of combination trials for EML4-ALK-driven lung tumors.

In general, the importance of fusion partners and/or the difference between variants of the same fusion partner in oncogenic fusions in lung cancer (e.g. *ROS1* and *RET* fusions) and across other cancer types (e.g. *NTRK*, *NRG1,* and *MET* fusions) remain largely unknown (Skoulidis and Heymach 2019; Consortium et al.). Our findings indicate that different variants of the same oncogenic fusion can be as functionally distinct from each other as distinct oncogenes. Consequently, beyond their role in driving expression and multimerization of kinase domains, fusion partners can profoundly shape the signaling landscape of tumors, thereby modulating the impact of other genomic alterations on tumorigenesis and potentially influencing treatment responses.

## METHODS

### Animal studies

All mouse experiments described in this study were approved by the Institutional Animal Care and Use Committee at Stanford University (protocol number 26696), or by the Regierungspräsidium Karlsruhe, Baden-Württemberg, Germany (protocol number G-265/19). Mice were kept in an environment free from specific pathogens, with a consistent light-dark cycle, and provided with standard mouse food and water *ad libitum*. *Kras^LSL-G12D/+^* (RRID:IMSR_JAX:008179), *R26^LSL-tdTomato^*(RRID:IMSR_JAX:007909), *H11^LSL-Cas9^* (RRID:IMSR_JAX:027632), *R26^LSL-Cas9-EGFP^* (RRID:IMSR_JAX:026179) and *Trp53*^flox/flox^ (RRID:IMSR_JAX:008462) mice have been previously described (Marino et al. 2000; Jackson et al. 2001; Madisen et al. 2010; Platt et al. 2014; Chiou et al. 2015).

### Analysis of human lung tumor data and tumor suppressor genes selection

The AACR Project Genomics, Evidence, Neoplasia, Information, Exchange (GENIE) database was used to select tumor suppressor genes associated with EML4-ALK lung cancer. Genomic alterations of EML4-ALK lung cancer patients were examined with a focus on identifying genomic mutations that provide critical insights into this specific cancer subtype and/or have been identified as significant regulators of lung cancer in previous research (**Supplementary Table 1**). GENIE 14.0-public data were collected through the Synapse platform (syn7844529, www.synapse.org). Patient samples were filtered to only include those annotated as the primary tumor, lung adenocarcinoma, and EML4-ALK positive as per structural variant annotations. Furthermore, samples were annotated according to their mutation status, categorized as wild-type, mutated, or not screened for each gene, considering the gene panels included in Project GENIE.

Genes that were mutated in at least 5 samples were considered. Additional genes of interest identified through a literature review were also included in the analysis (**Supplementary Table 1)**.

### Lentiviral vector generation

Triple sgRNA lentivectors were individually cloned and barcoded as previously using a modified protocol (Murray et al. 2019). sgRNAs targeting EGFP, intron 14 or 6 of *Eml4* (to generate the V1 or V3 inversion, respectively) and intron 19 of *Alk* ((Maddalo et al. 2014; Diaz-Jimenez et al. 2024), **Supplementary Figure S1h**, and **Supplementary Table 3**) were individually inserted downstream of the U6 promoters in the corresponding plasmids: pMJ114 (bovine U6; RRID:Addgene_85995), pMJ179 (mouse U6; RRID:Addgene_85996), pMJ117 (human U6; RRID:Addgene_85997), using site-directed mutagenesis (**Supplementary Figure S1h**). The U6-driven sgRNA cassettes were then PCR amplified to generate homology arms, which were placed in tandem using Gibson assembly into a PCR linearized Lenti-Cre vector (ref; RRID:Addgene_67594).

To mitigate the potential risk of intra-vector template-switching associated with the presence of duplicated DNA elements within the vector, we strategically employed a set of three distinct U6 promoters and three tracrRNAs, each sourced from different species and designed with nucleotide variations. Despite this, a notable degree of sequence similarity persisted.

Considering the potential for template switching between the U6 promoters and tracrRNAs, we systematically determined the optimal arrangement of sgRNAs within the triple vectors. This strategy favors template-switching events that remove either sg*Eml4* or sg*Alk*, leading to the production of lentiviral vectors that cannot initiate *Eml4-Alk* inversion and tumor formation.

The chosen configuration posed potential template switching events between specific elements, yielding distinct outcomes:

(1) tracrRNA#1 and tracrRNA#2: would remove sg*Eml4*, preventing the *Eml4-Alk* inversion and tumor development; (2) tracrRNA#2 and tracrRNA#3: would remove sg*Alk*, preventing the *Eml4-Alk* inversion and tumor development; (3) tracrRNA#1 and tracrRNA#3: would remove sg*Eml4* and sg*Alk*, preventing the *Eml4-Alk* inversion and tumor development; (4) U6#1 and U6#2: would remove sgTSG, allowing for the potential occurrence of *Eml4-Alk* inversion and tumor development, albeit with the inclusion of sgID for identifying sgTSG; (5) U6#2 and U6#3: would remove sg*Eml4*, preventing the *Eml4-Alk* inversion and tumor development; (6) U6#1 and U6#3: would remove sgTSG and sg*Eml4*, preventing the *Eml4-Alk* inversion and tumor development.

To target additional genes, sgEgfp was replaced with other sgRNAs in Lenti-sgEgfp-sgV1/Cre or Lenti-sgEgfp-sgV3/Cre through site-directed mutagenesis. This allowed for the targeting of specific tumor suppressor genes (Lenti-sgTSG-sgV1/Cre or Lenti-sgTSG-sgV3/Cre). Two to four distinct sgRNAs were used to target each putative tumor suppressor gene (**Supplementary Table 3**).

Lenti-sgTSG-sgV1/Cre or Lenti-sgTSG-sgV3/Cre vectors were diversified by the addition of sgID-BC as previously described ((Rogers et al. 2017; Murray et al. 2019); **Supplementary Table 3**). Distinct sgID-BCs flanked by BamHI and BspEI sites were generated by PCR amplification using the Lenti-Cre vector as a template, unique forward primers including the sgID-BC region, and a universal reverse primer. After amplification, the sgID-BC amplicons were digested with BamHI and BspEI and ligated into the corresponding linearized (BamHI and XmaI) Lenti-sgTSG-sgV1/Cre or Lenti-sgTSG-sgV3/Cre. The resulting diversified constructs were electroporated into bacteria and the bacterial colonies were counted and 90,000 to 5x10^5^ colonies were pooled. The plasmid DNA was isolated using PureLink HiPure Plasmid-Midiprep-Kit (Invitrogene, K210004). Because lentiviral template switching could generate integrated viral genomes with mismatches between sgRNAs and sgID-BCs (Hill et al. 2018; Li et al. 2021), we generated each viral vector separately to avoid template switching between vectors.

### Lentivirus production

Barcoded triple-sgRNA lentiviral/Cre vectors were produced as follows. Lenti-X 293T cells (Takara, #632180) were grown in DMEM with 10% FBS, 4 mM L-glutamine, and 1mM sodium pyruvate. Lenti-X 293T cells were co-transfected with packaging vectors (pCMV-dR8.2 RRID:Addgene_8455 and VSV-G RRID:Addgene_8454) using polyethyleneimine (Polysciences, #23966). 8 hours after transfection, cells were treated with 2.5 mM of sodium butyrate (Santa Cruz Biotechnology, #SC-202341B). Viral supernatant was collected at 48- and 72-hours post-transfection, concentrated by ultracentrifugation (1.5 h, 25,000 rpm, 4°C) and resuspended in PBS (Gibco, 10010023). The titer of the virus was determined as described previously (Rogers et al. 2018) using LSL-YFP mouse embryonic fibroblasts relative to a lab standard of known titer. Viral vectors were pooled and diluted in PBS to achieve the desired overall titer.

### Intratracheal delivery of lentivirus into the lung

8–14-week-old mice of both sexes were anesthetized by intraperitoneal injection of 100 µg/g ketamine and 14 µg/g xylazine. Tumors were initiated by intratracheal delivery of a maximum volume of 100 μl of the lentiviral preparations, as previously described (DuPage et al. 2009). The use of genetically modified organisms (GMO) was approved by the government of Baden-Württemberg, Germany (project number G-265/19, 81516 and 81520). To quantify the impact of inactivating different putative tumor suppressor genes, we initiated tumors with four different pools of barcoded Lenti-sgRNA/Cre vectors.

Injections were performed using **(1) Lenti-sg*TSG*^19^-sgV1/Cre** (**Figure 2, Supplementary Figure S2,S3**). A titer of 2e6 IFU was injected into 13 *R26^LSL-Cas9-GFP/LSL-Cas9-GFP^* (*Cas9-EGFP*) and a titer of 1.5e5 IFU was injected into 1 *Kras^LSL-G12D/+^;p53^flox/wt^*and 5 *Kras^LSL-G12D/+^;p53^wt/wt^* (collectively Cas9-negative control) mice. **(2) Lenti-sg*TS1*^19^-sgV*3*/Cre** (**Figure 2, Supplementary Figure S2,S3**). A titer of 2e6 IFU was injected into 14 *R26^LSL-Cas9-GFP/LSL-Cas9-GFP^*(*Cas9-EGFP*) and a titer of 1.5e5 IFU was injected into 3 *Kras^LSL-G12D/+^;p53^flox/wt^*and 3 *Kras^LSL-G12D/+^;p53^wt/wt^* (collectively Cas9-negative control) mice. **(3) Lenti-sg*TSG*^75^-sgV1/Cre** (**Figure 3, Supplementary Figure S4,S5**). A titer of 8.7e5 IFU was injected into 22 *R26^LSL-Cas9-GFP/LSL-Cas9-GFP^* (*Cas9-EGFP*) and a titer of 1e5 IFU was injected into 5 *Kras^LSL-G12D/+^;p53^flox/wt^* (Cas9-negative control) mice. **(4) Lenti-sg*TSG*^75^-sgV3/Cre** (**Figure 3, Supplementary Figure S4,S5**). A titer of 7.5e5 IFU was injected into 27 *R26^LSL-Cas9-GFP/LSL-Cas9-GFP^* (*Cas9-EGFP*) and a titer of 1e5 IFU was injected into 6 *Kras^LSL-G12D/+^;p53^flox/flox^* (Cas9-negative control) mice.

### Histology and immunohistochemistry

Lung tissues were perfused with PBS through the heart and fixed overnight in 10% formalin (Sigma, HT501128), transferred to 50% ethanol, processed in a tissue processor (Leica ASP300S), and embedded in paraffin blocks. Sections were cut at 3-µm thickness and stained for hematoxylin and eosin (H&E) for histologic examination. For IHC, slides were dewaxed, and antigen retrieval was performed using a citrate-based solution pH 6.0 (Vector Laboratories, H-3300). Endogenous peroxidase was blocked using 3% H2O2, and endogenous species protein were blocked using the appropriate species serum depending on the secondary antibody. Tissues were incubated with primary antibodies overnight. Primary antibodies used were: Rabbit Anti-Prosurfactant Protein C (Millipore Cat#AB3786, RRID:AB_91588), Rabbit Anti-Clara Cell Secretory Protein (Millipore Cat#07-623, RRID:AB_310759), Rabbit Anti-TTF1 (Abcam Cat#ab76013,RRID:AB_1310784), Rabbit Anti-GFP (Cell Signaling Technology Cat#2956, RRID:AB_1196615), and Rabbit Anti-H3K36me3 (Cell Signaling Cat#4909, RRID:AB_1950412,). The ABC kit (Vector Laboratories, PK-6101) and DAB peroxidase substrate kit (Vector Laboratories, SK-4100) were used for signal detection and tissues were counterstained with hematoxylin. Pictures were taken in a Tissuegnostic TissueFAX system. Histological analyses were performed using StrataQuest software (TissueGnostics) or QuPath (version 0.4.4).

### Dissociation of individual microdissected tumors

Individual microdissected tumors were minced into <0.5 mm^3^ pieces and transferred into pre-warmed digestion media (Dispase (5 U/mL; Corning #354235), DNase I, Grade II (1 mg/mL; Sigma #10104159001), Y-27632 ROCK inhibitor (0.5 mM; Stem Cell Technologies #72304) in Advanced DMEM/F-12 (Gibco #12634010)). The minced tumors were digested at 37°C for 40 minutes with gentle horizontal shaking. Tumors were vigorously disrupted by pipetting every 15 minutes. The digestion media was then neutralized with stop media (DMEM/F-12 (VWR International #392-0412), FBS (10%, Gibco #10270-106), HEPES (0.01 M, Gibco #15630056) with DNase I, Grade II (0.1 mg/mL), and ROCK inhibitor (0.05 mM)) and cell suspensions were passed through 40 µm cell strainers. The samples were centrifuged at 1,100 rpm for 7 minutes and pellets were resuspended in red blood cell lysis buffer (Sigma-Aldrich, R7757). Cells were washed with FACS buffer by spinning at 1,100 rpm for 7 minutes. Cell pellets were then resuspended in an appropriate volume of FACS buffer.

### Fluorescence-Activated Cell Sorting (FACS)

Single-cell suspensions from individual tumors were prepared as described above. Cells were stained with Fc block and an antibody cocktail (APC anti-mouse CD45 (BioLegend Cat#103112 RRID:AB_312977), APC anti-mouse TER-119/Erythroid (CellsBioLegend #116212, RRID:AB_313713), APC anti-mouse CD31 (BioLegend #102409, RRID:AB_312904), APC anti-mouse/human CD11b (BioLegend #101211, RRID:AB_312794), DAPI solution (Life Technologies #62248))

Cell suspensions were incubated for 30 minutes on ice and washed twice with FACS buffer by centrifuging the samples at 1,200 rpm for 5 minutes at 4°C. Prior to cell sorting, DAPI (Thermo Scientific, 62248) was added at a final concentration of 5 µg/mL. 4-Way Purity FACS was performed using a BD FACS Aria or Fusion sorter. Cancer cells were sorted as DAPI^neg^/CD45^neg^/CD31^neg^/CD11b^neg^/TER-119^neg^/GFP^pos^.

### DNA extraction and tumor genotyping

Genomic DNA from sorted tumor cells was extracted using the QIAamp DNA Mini Kit (Qiagen, #51304) according to the manufacturing protocol. Purified DNA was then genotyped by PCR using the HotStart Master Mix Kit (Qiagen, #203443). Amplicons were then sequenced by Sanger sequencing through Mycrosynth services. The Sanger products’ DNA were then examined using Benchling [Biology Software] (2018-2024, https://benchling.com). TIDE was used to estimate the spectrum and frequency of small insertions and deletions using wild-type sequences as references.

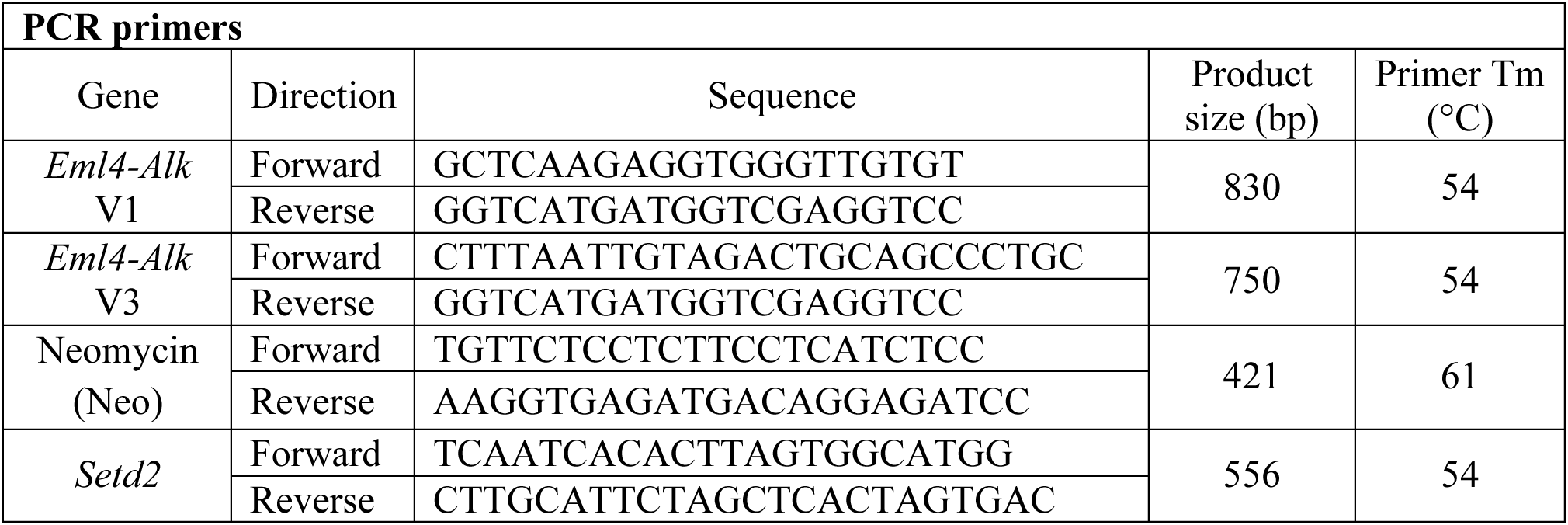

### Library preparation for Tuba-Seq

Genomic DNA was isolated from bulk tumor-bearing lung tissue from each mouse as previously described (Cai et al. 2021). Briefly, benchmark control cell lines were generated from LSL-YFP MEFs transduced by a barcoded Lenti-sgNT3/Cre vector (NT3: an inert sgRNA with a distinct identifying sequence or ‘sgID’) and purified by sorting YFP^pos^ cells. Three benchmark control cell lines (500,000 cells each) were added to each mouse lung sample prior to lysis to enable the calculation of the absolute number of neoplastic cells in each tumor from the number of sgID-BC reads. Following homogenization and overnight protease K digestion, genomic DNA was extracted from the lung lysates using standard phenol-chloroform and ethanol precipitation methods. Subsequently, Q5 High-Fidelity 2x Master Mix (New England Biolabs, M0494X) was used to amplify the sgID-BC region from 32 μg of genomic DNA in a total reaction volume of 800 μl per sample. The unique dual-indexed primers used were Forward: AAT GAT ACG GCG ACC ACC GAG ATC TAC AC-8 nucleotides for i5 index-ACA CTC TTT CCC TAC ACG ACG CTC TTC CGA TCT-6 to 9 random nucleotides for increased diversity-GCG CAC GTC TGC CGC GCT G and Reverse: CAA GCA GAA GAC GGC ATA CGA GAT-6 nucleotides for i7 index-GTG ACT GGA GTT CAG ACG TGT GCT CTT CCG ATC T-9 to 6 random nucleotides for increased diversity-CAG GTT CTT GCG AAC CTC AT. The PCR products were purified with Agencourt AMPure XP beads (Beckman Coulter, A63881) using a double size selection protocol. The concentration and quality of the purified libraries were determined using the Agilent High Sensitivity DNA kit (Agilent Technologies, 5067-4626) on the Agilent 2100 Bioanalyzer (Agilent Technologies, G2939BA). The libraries were pooled based on lung weight to ensure even sequencing depth, cleaned up again using AMPure XP beads and sequenced (read length 2x150bp) on the Illumina HiSeq 2500 or NextSeq 500 platform (Admera Health Biopharma Services).

### Analysis of sgID-BC sequencing data

Sequencing of Tuba-seq libraries produces reads that are expected to contain an 8-nucleotide sgID followed by a 30-nucleotide BC of the form GCNNNNNTANNNNNGCNNNNNTANNNNNGC, where each of the 20 Ns represents a random nucleotide. Each sgID has a one-to-one correspondence with an sgRNA in the viral pool (**Supplementary Table 3**); thus, the sgID sequence identifies the gene targeted in each tumor. Note that all sgID sequences in a viral pool differ from each other by at least three nucleotides such that incorrect sgID assignment (and thus, inference of tumor genotype) due to PCR or sequencing error is extremely unlikely. Distinct sets of sgIDs were used to tag each sgRNA in the V1 and V3 viral pools. The random 20-nucleotide BC tags all cells in a single clonal expansion. Note that the length of the BC ensures a high theoretical potential diversity (∼4^20^ > 10^12^ BCs per vector), with the actual diversity of each Lenti-sgRNA/Cre vector dictated by the number of bacterial colonies pooled during the plasmid barcoding step.

FASTQ files were parsed using regular expressions to identify the sgID and BC in each read. To minimize the effects of sequencing error on BC identification, we required the forward and reverse reads to agree completely within the 30-nucleotide sgID-BC region. We also performed an analysis to identify BCs that were likely to have arisen from genuine tumors due to PCR or sequencing errors. Given the low rate of sequencing error, we expect these ‘spurious tumors’ to have read counts that are far lower than the read counts of the genuine tumors from which they arise. To minimize the impact of these ‘spurious tumors’, we identified small ‘tumors’ with BCs that were highly similar to the BCs of larger tumors in the same sample. Specifically, if a pair of ‘tumors’ within a sample had BCs that were within a Hamming distance of two, and if one of the tumors had fewer than 5% as many reads as the other, then the reads associated with the smaller tumor were attributed to the larger tumor.

After these filtering steps, the read counts associated with each BC were converted to absolute neoplastic cell numbers by normalizing to the number of reads from the ‘spike-in’ cell lines added to each sample before lung lysis and DNA extraction.

### Removal of contaminating barcodes

sgID-BC sequences that are not from genuine tumors in an individual tumor-bearing lung sample can none-the-less be present in sequencing libraries for several reasons including intra-experiment sample-to-sample cross-contamination during library preparation, external contamination (*e.g*., from samples and libraries from other experiments), and library-to-library misassignment during sequencing. These sgID-BC sequences have the potential to be identified as small tumors (*i.e*., ‘spurious tumors’) and thereby reduce the precision of our analyses. Both external contamination and sample-to-sample contamination result in the identification of the same sgID-BC in multiple samples. However, some sgID-BCs are expected to recur across samples in the absence of contamination due to the finite diversity of the sgID-BC region in each lenti-sgRNA/Cre vector. To minimize the effects of contamination, we examined patterns of sgID-BC recurrence across samples and removed barcodes that occurred in a number of samples that would be highly unlikely to occur by chance given the BC diversity of each vector.

To estimate the BC diversity associated with each sgID, we assume that the probability of observing a BC in *i* mice is Poisson distributed: P(*k*=*i*; λ) = *λ*^k^ *e*^−*λ*^ */ k*!, where *λ* is the mean number of mice that barcodes appear in for a given lenti-sgRNA vector. To estimate *λ*_r_ for each sgID *r* in the dataset we note that *λ*_r_/(1 – *e^−λ^*^r^) = *μ*_non-zero_, where *μ*_non-zero_ = Σ ^∞^P(*k=i*; *λ*_r_) is the mean number of occurrences of each BC that occurred once or more (a known quantity). Given the Poisson distribution defined for each sgID *r*, we calculated a lenti-sgRNA/Cre vector specific threshold N_r_ such that 99.9% of barcodes would be expected to appear in N_r_ samples or fewer and identified and removed all ‘tumors’ with sgID-BCs that occurred in a number of samples exceeding N_r_.

sgID-BCs can also recur across samples due to misassignment of reads during sample de-multiplexing. While misassignment of reads is expected to be extremely rare, it could result in sgID-BCs from genuine tumors with very large read counts in one sample (i.e., a very large, genuine tumor) appearing with low read counts in additional samples sequenced in the same lane. To guard against this possibility and ensure that very large tumors were not systematically discarded by the procedure described in the prior paragraph, we examined the distribution of reads across samples to identify sgID-BCs that appeared in >N_r_ samples but where >95% of sequencing reads were assigned to a single sample. These sgID-BCs were retained in the sample with the bulk of the sequencing reads and discarded in all other samples. All other barcodes appearing in > N_r_ samples were discarded from all samples.

### Removal of mice with low tumor number

After processing the sgID-BC sequencing data and performing the filtering steps described above the number of unique barcodes represents the number of genuine clonal tumors. Tumor number across mice was highly variable in all cohorts (i.e., the mean number of tumors in mice transduced with the Lenti-sg*TSG*^75^-sg*V1*/Cre pool was 4,159 with a standard deviation of 3,553 and the mean number of tumors in mice transduced with the Lenti-sg*TSG*^75^-sg*V3*/Cre pool was 9,403 with a standard deviation of 6,310). This is most likely driven by variability in lentiviral delivery and/or other factors that change overall transduction across mice. Samples with extremely low tumor number (fewer than 1,000 tumors detected) were deemed to not have been successfully transduced and were removed from the analysis. We removed one sample from the *Cas9-EGFP* cohort transduced with Lenti-sg*TSG*^19^-sg*V3*/Cre, four samples from the *Cas9-EGFP* cohort transduced with Lenti-sg*TSG*^75^-sg*V1*/Cre, one sample from the *Cas9-EGFP* cohort transduced with Lenti-sg*TSG*^75^-sg*V3*/Cre, and one sample from the Cas9-negative *Kras^G12D^;Trp53^flox/wt^* cohort transduced with Lenti-sg*TSG*^75^-sg*V1*/Cre.

### Filtering of vectors with insufficient titer from analysis

Each Lenti-sgRNA/Cre vector was titered and pooled with the goal of equal representation of each vector in the viral pool, however, our analysis requires us to know the exact proportion of each vector in the pool. We therefore transduced cohorts of Cas9-negative control mice with each lentiviral pool to quantify the representation of each vector in each pool. These cohorts were composed of *Kras^LSL-G12D/+^;p53^flox/wt^*and *Kras^LSL-G12D/+^; p53^flox/flox^* mice which do not express Cas9. Cells transduced in these mice will still expand into *Kras*-driven tumors with unique sgID-BCs, but all sgRNAs are functionally inert. As a result, the relative tumor number associated with each sgRNA in these mice reflects the proportion of that vector in that viral pool. Quantifying the number of tumors associated with each sgRNA in these Cas9-negative mice revealed a small number of vectors that were represented at too low a proportion to enable further analysis. Vectors with an average of fewer than 15 unique barcodes per mouse in the Cas9-negative control cohorts were removed prior to analysis of tumor suppressive effects (*sgDnmt3a#1*, *sgp53#1*, *Nkx2-1#1 and sgKmt2c#1* were removed from analysis of the Lenti-sg*TSG*^75^-sg*V1*/Cre viral pool; *sgSmad4#2*, *sgSlc34a2#2* and *sgLats2#1* were removed from the analysis of the Lenti-sg*TSG*^75^-sg*V3*/Cre pools. No vectors were removed from analyses of the Lenti-sg*TSG*^19^-sg*V1*/Cre and Lenti-sg*TSG*^19^-sg*V3*/Cre viral pools).

### Adaptive sampling of tumors for statistical comparison of tumor genotypes

We performed four independent screens of tumor suppressor function in EML4-ALK-driven lung cancer using different viral pools (Lenti-sg*TSG*^19^-sg*V1*/Cre, Lenti-sg*TSG*^19^-sg*V3*/Cre, Lenti-sg*TSG*^75^-sg*V1*/Cre, and Lenti-sg*TSG*^75^-sg*V3*/Cre). These experiments varied in several respects including technical differences such as sequencing depth, cohort size, and the titer and pooling of Lenti-sgRNA/Cre vectors, as well as intrinsic differences in the growth dynamics of V1- and V3-driven tumors. To mitigate the impact of these factors and ensure that an equivalent portion of the tumor size distribution was analyzed for each sgRNA 𝑖 in each experiment 𝑗, we scaled the number of tumors analyzed from each dataset (combination of sgRNA and experiment) to account for differences in viral titer and pooling and then analyzed the largest 𝑁*_i,j_* tumors per vector.

This scaling procedure requires selecting a benchmark (basal) sgRNA and a benchmark (basal) experiment, and then selecting a defined number of tumors with that sgRNA from that experiment (𝑁*_i=basal,j=basal_*). The number of tumors sampled for each other sgRNA 𝑖 in each experiment 𝑗 (𝑁*_i,j_*) is then adjusted to take into account differences in both the overall viral titer delivered and in the representation of sgRNAs between the viral pool used in experiment j and the viral pool used in the basal experiment:

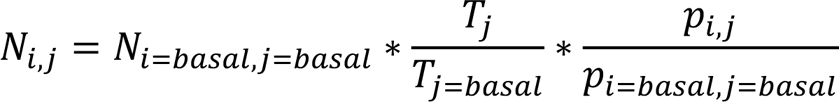

where 𝑇_j_ denotes the total viral titer delivered in experiment *j* and 𝑝*_i,j_* denotes the proportion of the viral pool in experiment 𝑗 allocated to sgRNA 𝑖. 𝑇_j_is a known quantity determined during viral preparation which represents the sum of the viral titers delivered to each mouse in experiment 𝑗. 𝑃*_i,j_* values were calculated using Tuba-seq data generated by transducing control *Kras^LSL-G12D/+^;R26^LSL-Tom^;p53^flox/wt^*and *Kras^LSL-G12D/+^;R26^LSL-Tom^;p53^flox/flox^*mice. As these mice lack Cas9 expression all Lenti-sgRNA/Cre vectors are functionally inert, and the observed tumor number associated with each sgRNA reflects the make-up of the viral pool. To account for variation in tumor number across mice, the sample of 𝑁_*i,j*_tumors was distributed across all mice in the cohort in proportion to the total number of tumors in each mouse.

For analyses of data generated using the Lenti-sg*TSG*^19^-sg*V1*/Cre and Lenti-sg*TSG*^19^-sg*V3*/Cre viral pools (**Figure 2**), we used sgNT1 (a non-targeting control sgRNA) in the V1 cohort as the basal dataset. Likewise, for analyses of data generated using the Lenti-sg*TSG*^75^-sg*V1*/Cre, and Lenti-sg*TSG*^75^-sg*V3*/Cre viral pools (**Figures 3** and **5**) we used sgNT1 in the V1 cohort as the basal dataset. Note that these choices were arbitrary and identical results could be achieved by basing the sampling calculations on any sgRNA in any dataset.

**^Selection of^** ^𝑁^*_i=basal,j=basal_*

The parameter 𝑁*_i=basal,j=basal_* defines the portion of the tumor size distribution that we use in our calculations of tumor suppressive effects. In selecting 𝑁*_i=basal,j=basal_* for each analysis, we sought to include as much data as possible while maintaining high data quality by excluding tumors with barcodes supported by very few reads. We sequenced samples to median depths of 35.2, 44.8, 15.5, and 16.0 cells per read for the cohorts transduced with Lenti-*sgTSG^19^-sgV1/Cre*, Lenti-*sgTSG^19^-sgV3/Cre*, Lenti-*sgTSG^75^-sgV1/Cre*, and Lenti-*sgTSG^75^-sgV3/Cre*, respectively. We reasoned that restricting our analysis to tumors above 250 cells in size (and therefore supported by an average of >5 reads in our lowest depth dataset) would minimize noise associated with the detection and measurement of very small tumors while maintaining high statistical power to detect tumor suppressive effects. We therefore selected an 𝑁*_i=basal,j=basal_* for each analysis that corresponded to the number of tumors with >250 cells for the basal sgRNA in the basal experiment. The values of 𝑁*_i=basal,j=basal_*used for each experiment are shown in the top row of the table below. To ensure that our findings were robust to a range of minimum tumor sizes and corresponding numbers of tumors sampled, we also performed our analyses using values of 𝑁*_i=basal,j=basal_* corresponding to the inclusion of tumors larger than 300, 350, and 400 cells and found similar results (**Supplementary Figure S8c**).

| | Number of sgNT1 tumors sampled in V1 cohort ($N_{i=basal,j=basal}$)<br>rounded to the nearest hundred | |
| --- | --- | --- |
| Minimum size Threshold | Lenti-*sgTSG<sup>19</sup>-sgV1/Cre* and Lenti-*sgTSG<sup>19</sup>-sgV3/Cre* experiment<br>(**Figure 2**) | Lenti-*sgTSG<sup>75</sup>-sgV1/Cre*, and Lenti-*sgTSG<sup>75</sup>-sgV3/Cre* (**Figures 3 and 5**) |
| 250 | 700 | 700 |
| 300 | 600 | 700 |
| 350 | 500 | 600 |
| 400 | 400 | 600 |

The same logic was used to determine 𝑁*_i=basal,j=basal_* for the control Cas9-negative cohorts in each experiment. Because these cohorts contained both *Kras^LSL-G12D/+^;R26^LSL-Tom^;p53^flox/wt^*and *Kras^LSL-G12D/+^;R26^LSL-Tom^;p53^flox/flox^* mice, a given lentiviral titer may not initiate the same number of tumors in each cohort (although the proportion of each sgRNA should be the same as there is no Cas9). We therefore defined separate 𝑁*_i=basal,j=basal_* values for each control cohort (again using sgNT1 as our basal sgRNA) and performed the scaling calculation described above within each cohort. The 𝑁*_i=basal,j=basal_* sampled for these cohorts are shown in the table below.

| Minimum<br>size<br>Threshold | $N_{i=basal,j=basal}$<br>Lenti-sgTSG<sup>19</sup>-<br>sgV1/Cre<br>control cohort | $N_{i=basal,j=basal}$<br>Lenti-sgTSG<sup>19</sup>-<br>sgV3/Cre<br>control cohort | $N_{i=basal,j=basal}$<br>Lenti-sgTSG<sup>75</sup>-<br>sgV1/Cre control<br>cohort | $N_{i=basal,j=basal}$<br>Lenti-sgTSG<sup>75</sup>-<br>sgV3/Cre<br>control cohort |
| --- | --- | --- | --- | --- |
| 250 | 1000 | 1100 | 100 | 400 |
| 300 | 900 | 1000 | 100 | 300 |
| 350 | 900 | 900 | 100 | 300 |
| 400 | 800 | 800 | 100 | 200 |

### Summary statistics for the impact of gene inactivation on tumorigenesis

After performing the adaptive sampling described above, we assessed the extent to which each targeted gene (*X)* affects tumor initiation and growth, by comparing the distribution of tumor sizes produced by vectors targeting that gene (sg*X* tumors) to the distribution produced by our negative control vectors (sg*Inert* tumors). We relied on two statistics to characterize these distributions: the log-normal mean size (adaptively sampled LN mean or ASM) and the size of tumors at the 95^th^ percentile of the distribution (adaptively sampled 95^th^ percentile size). The LN mean is the maximum-likelihood estimate of mean tumor size assuming a log-normal distribution. Previous work found that this statistic represents the best parametric summary of tumor growth based on the maximum likelihood quality of fit of various common parametric distributions (Rogers et al. 2017). The 95^th^ percentile size is a nonparametric summary statistic of the tumor size distribution. This metric corresponds to the right tail of the distribution and thus reflects the growth of larger tumors, thereby avoiding issues stemming from potential variation in cutting efficiency among guides.

Note that while these statistics are functions of the tumor size distribution, they are also sensitive to the effects of tumor suppressor gene inactivation on tumor initiation due to our use of adaptive sampling. For example, a gene that increases tumor burden when inactivated solely by increasing the number of tumors (i.e., without shifting the tumor size distribution) would still increase the LN mean and 95^th^ percentile size of the adaptively sampled datasets and would therefore still be detectable as a tumor suppressor using these metrics. These statistics therefore integrate the impact of each targeted gene on the initiation and growth of tumors.

To quantify the extent to which each gene suppressed or promoted tumorigenesis, we normalized statistics calculated on tumors of each genotype to the corresponding inert statistic. The resulting ratios reflect the fitness advantage (or disadvantage) associated with each tumor genotype relative to the initiation and growth of sg*Inert* tumors.

The adaptively sampled relative 95^th^ percentile size for tumors of genotype X was calculated as:

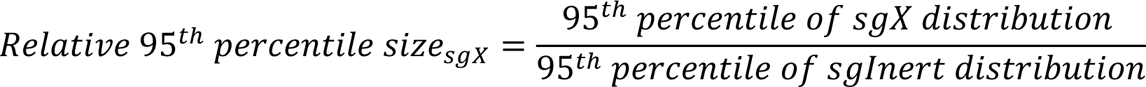

Likewise, the adaptively sampled relative LN mean size for tumors of genotype X was calculated as:

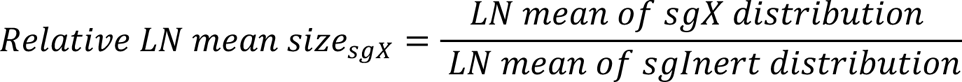

For both statistics the sg*X* and sg*Inert* distributions were adaptively sampled as previously described and the sg*Inert* statistic is the median value across all inert vectors.

### Aggregating information across sgRNAs in the calculation of tumor growth metrics

Adaptive sampling and calculation of summary statistics were performed per sgRNA. To calculate statistics at the gene level, we aggregated information across all sgRNAs targeting the same gene. Specifically, the gene-level statistics reported are the weighted average of the per-sgRNA statistics, where the contribution of each sgRNA was proportional to the number of adaptively sampled tumors with that sgRNA.

### Calculation of confidence intervals and P-values for the impacts of gene inactivation on tumorigenesis

Confidence intervals and P-values were calculated using bootstrap resampling to estimate the sampling distribution of each statistic. To account for both mouse-to-mouse variability and variability in tumor size and number within mice, we adopted a two-step, nested bootstrap approach where we first resampled mice, and then resampled tumors within each mouse in the pseudo-dataset. 10,000 bootstrap samples were drawn for all reported P-values. 95% confidence intervals were calculated using the 2.5th and 97.5th percentiles of the bootstrapped statistics. Because we calculate metrics of tumor growth that are normalized to the same metrics in sg*Inert*

tumors, under the null model where genotype does not affect tumor growth, the test statistic is equal to 1. Two-sided p-values were thus calculated as follows:

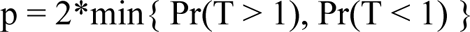

Where T is the test statistic and Pr(*T*>1) and Pr(*T*<1) were calculated empirically as the proportion of bootstrapped statistics that were more extreme than the baseline of 1. To account for multiple hypothesis testing, p-values were FDR-adjusted using the Benjamin-Hochberg procedure as implemented in the Python package stats models. Summarized statistics of all Tuba-seq experiments in this study can be found in **Supplementary Tables 2,4**.

### Calculation of p-values for differential effects of tumor suppressor inactivation in EML4-ALK V1 and V3 tumors

To compare the impacts of inactivating a given tumor suppressor gene between tumors driven by EML4-ALK V1 and EML4-ALK V3, we defined a test statistic T equal to the difference between the effect (either magnitude of effect or rank-order of magnitude) in V1-driven and V3-driven tumors. We produced a bootstrapped distribution of T by resampling mice and tumors within each cohort (as described above) and repeatedly calculating T. Under the null hypothesis, the impact of inactivating a given tumor suppressor gene is the same in V1-driven and V3-driven tumors and T is therefore equal to 0. Two-sided p-values were thus calculated as followed:

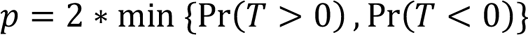

Where Pr(T>0) and Pr(T<0) were calculated empirically as the proportion of bootstrapped statistics that were more extreme than the baseline of 0. To account for multiple hypothesis testing, p-values were FDR-adjusted using the Benjamini-Hochberg procedure as implemented in the Python package stats models or in R Stats package.

### Analyses of correlation in tumor suppressive effects across oncogenic contexts

To assess the extent to which the effects of tumor suppressor gene inactivation were similar across oncogenic contexts, we calculated Spearman’s rank correlation coefficient (ρ) of the effects of tumor suppressor inactivation between pairs of oncogenic contexts.

For comparison of our EML4-ALK V1 and V3 datasets (both V1 vs V3, and “self” comparisons of V1 vs V1 and V3 vs V3), we calculated ρ using the adaptively sampled log-normal mean tumor sizes (ASMs) for each putative tumor suppressor in each dataset. Confidence intervals around the point estimate of ρ were generated by bootstrapping each dataset using a two-step nested resampling approach (first resampling mice, then resampling tumors within each bootstrapped mouse), calculating the ASM for each gene in each pseudo-dataset, and then calculating ρ for the bootstrapped ASMs. 10,000 bootstrap samples were drawn and 95% confidence intervals were calculated using the 2.5^th^ and 97.5^th^ percentiles of the bootstrapped statistics.

For comparisons using data from (Blair et al. 2023) and the independently generated KRAS^G12D^ dataset (both cross-oncogene and “self” comparisons), ρ was calculated using the relative 95^th^ percentile tumor size for each gene. Confidence intervals were generated by fitting normal distributions to the reported 95% confidence interval around the relative 95^th^ percentile tumor size for each gene, repeatedly sampling statistics for each gene from these distributions, and then calculating ρ using the resampled statistics. 10,000 samples were drawn and 95% confidence intervals were calculated using the 2.5^th^ and 97.5^th^ percentiles of the resampled statistics.

For each set of tumor suppressor genes, we calculated the distribution of ρ under the null hypothesis (that there is no correlation in effects between two contexts) using a simulation that repeatedly measured the correlation between the list of genes and a randomly shuffled version of the list. The simulation was run 10,000 times for each gene set, and 95% confidence intervals were calculated using the 2.5^th^ and 97.5^th^ percentiles of the simulated statistics.

### Investigation of an independent real-world NSCLC cohort

In this study, we performed a retrospective analysis of 95,641 patients with non-small cell lung cancer (NSCLC), who received tissue biopsy-based comprehensive genomic profiling (CGP) from August 2014 through March 2023, using FoundationOne^®^ or FoundationOne^®^CDx, during routine clinical care. DNA was extracted from formalin-fixed paraffin-embedded tissue sections and hybrid capture was performed on up to 324 cancer-associated genes, including select introns from genes frequently rearranged in cancer, to detect different classes of genomic alterations: short variants (base substitutions, short insertions/deletions), gene copy number amplifications and homozygous deletions, as well as gene fusions and rearrangements. Details on the sequencing and the analytical validation of the assays have been described previously (Frampton et al. 2013; Milbury et al. 2022).

Specifically, among cases with a detected EML4-ALK fusion, the breakpoint information was used to classify the different variants: V1 (Intron 19 of ALK/Intron 13 of EML4), V2 (Intron 19 of ALK/Intron 20 of EML4), V3 (Intron 19 of ALK/Intron 6 of EML4), V5 (Intron 19 of ALK/Intron 2 of EML4) and V5’ (Intron 19 of ALK/Intron 18 of EML4), and other (capturing other EML4-ALK fusions). Rare cases with multiple detected EML4-ALK variants were excluded from the analysis. The spectrum and prevalence of co-occurring alterations were assessed for cases with different EML4-ALK variants; the 296 genes commonly targeted on both FoundationOne^®^ or FoundationOne^®^CDx were examined for these analyses. A Fisher’s exact test was used to test for difference in the prevalence of co-occurring gene alterations between two different EML4-ALK variant groups. Two-sided p-values were calculated, followed by adjustment for multiple testing using the Benjamini–Hochberg false discovery rate (FDR) method; an FDR-adjusted p-value ≤ 0.05 was considered statistically significant. Statistics, computation, and plotting were carried out using Python 3.9.12 (Python Software Foundation) and R 4.3.1 (R Foundation for Statistical Computing).

Approval for the study of this cohort, including a waiver of informed consent and Health Insurance Portability and Accountability Act waiver of authorization, was obtained from the Western Institutional Review Board (Protocol #20152817).

### Power calculation to assess ability to detect differences in mutation frequency between V1- and V3-driven tumors

Power calculations were performed to evaluate the ability to detect differences in mutation frequency between V1- and V3-driven tumors given the 1) the number of samples with V1 and V3 fusions in the EML4-ALK NSCLC cohort, 2) variable overall frequencies of alteration, and 3) variable relative frequencies of alteration in V1 vs V3 samples. Simulations were performed using the power.fisher.test function implemented in the R package statmod, which assumes a binomial distribution for mutation counts for each cohort and calculates a two-sided Fisher’s exact test. 10,000 simulations were performed for each parameter combination.

### RNA extraction and RNA library preparation from sorted tumor cell genotyping

RNA from sorted cells was isolated using the PicoPure RNA Isolation Kit (Life Technologies, #KIT0204) according to the manufacturer’s instructions. Subsequently, RNA-sequencing libraries were generated using the SMART-Seq mRNA library prep (Takara, #634768) and Unique Dual Index Kit (Takara, #634756) following manufacturer’s protocol. Input included 1 ng of RNA and 1 ng of cDNA. Quality control was performed by assessing RNA, cDNA, and final library qualities using Tapestation. Libraries were sequenced in a NovaSeq 6k PE 150 S4.

### RNA sequencing analysis

Samples were aligned to the mouse genome GRCm38mm10 using STAR (Version 2.5.3a). Downstream analysis was done using R statistical computing software (Version 4.2). Counts were processed using DESeq2 (Love et al. 2014) (Version 1.38.3). Sex of the mice were included in the design formula as covariable. The data was normalized using Variance stabilizing transformation (VST), and differential expression analysis was carried out with DESeq2. PCA was computed using the top 6000 most variable genes using package FactoMineR (Lê et al. 2008), and visualization using the package Factoextra. Pathway activities were calculated using decoupleR (Badia-i-Mompel et al. 2022).

### Data availability statement

All barcode sequencing datasets will be made publicly available through the NCBI Sequence Read Archive Database and accession numbers provided before the publication of the manuscript. All single-cell RNA sequencing data will be made publicly available through the Gene Expression Omnibus and accession numbers provided before the publication of the manuscript.

For the Foundation Medicine cohort, the data were derived from clinical samples. Data supporting the findings within this cohort have been provided within the article and its supplementary files. Due to HIPAA requirements, we are not authorized to share underlying sequence data or individualized patient genomic data, which contain potentially identifying or sensitive patient information. Foundation Medicine, Inc. is committed to collaborative data analysis, and has well-established and widely utilized mechanisms by which investigators can query its genomic database of >800,000 deidentified sequenced cancers to obtain aggregated datasets. More information and mechanisms for data access can be obtained by contacting the corresponding author or the Foundation Medicine Data Governance Council at.

## Acknowledgments

We thank the DKFZ Core Facilities for Light Microscopy and Flow Cytometry for their outstanding technical support, the Central Animal Facility for their excellent care of the animals, and members of the Sotillo, Petrov, and Winslow laboratories for helpful comments. A.D.J. was supported by the DKFZ International PhD Program and by the DZL Academy Mobility Grant. E.G.S was supported by a Tobacco-Related Disease Research Program (TRDRP) predoctoral fellowship (T33DT6556). C.W.M. was supported by the NSF Graduate Research Fellowship Program and an Anne T. and Robert M. Bass Stanford Graduate Fellowship. E.G.S. was supported by a TRDRP Predoctoral Fellowship (T33DT6556). This work was supported by the Worldwide Cancer Research Grant 22-0197, the German Center for Lung Research #82DZL004B4 (to R.S.), and NIH R01-CA234349 (to M.M.W. and D.A.P.).

## Author contributions

Conceptualization and design of the study: A.D.J., E.G.S., M.M.W., R.S.

Designed and constructed all DNA vectors: A.D.J., K.S. with contributions from C.W.M.

Conducted all autochthonous lung modeling: A.D.J., M.K.T.

Performed histology stains: A.D.J., F.A.

Prepared Tuba-seq libraries from mouse lung: A.D.J., L.A.

Designed the computational pipeline, methodology, and formal analysis: E.G.S., D.A.P.

Analysis of human tumor sequencing data: S.S., S.D.S, E.G.S, E.S.S.

RNA-seq analysis: A.D.J.; O.G., B.B.

Wrote the manuscript: A.D.J., E.G.S., M.M.W., R.S.

Resources: A.D.J., M.M.W., R.S.

---

**Supplementary Figure 1.**
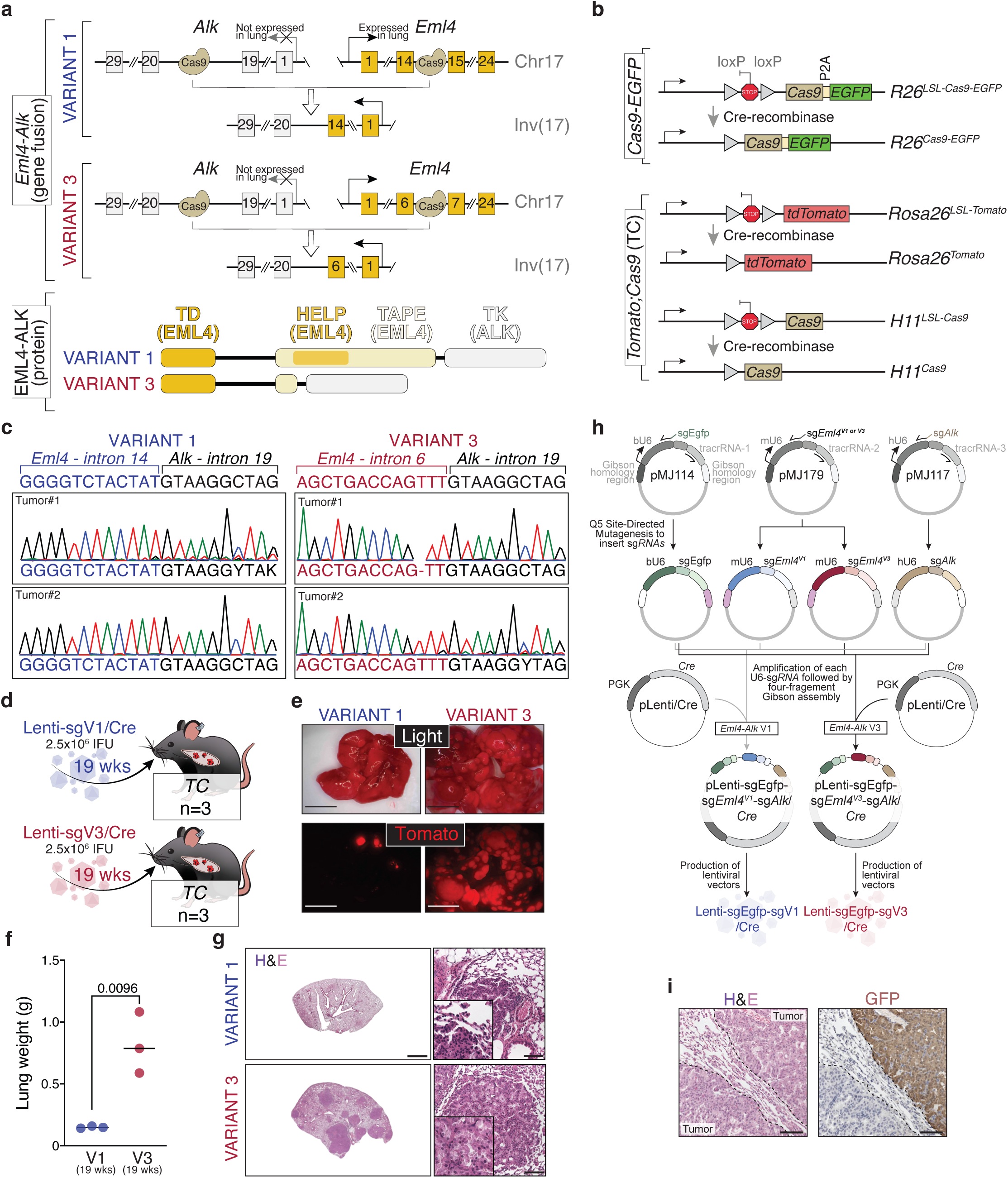
Lentiviral induced EML4-ALK V1 and V3 tumors in *R26^LSL-Cas9-EGFP^* mice and targeting of EGFP. **a.** Upper panel shows the exon structure of *Eml4*, *Alk*, and the *Eml4-Alk* V1 and V3 fusion variants, with squares indicating exons. The lower panel shows the protein products resulting from the *Eml4-Alk* chromosomal rearrangement, highlighting distinctive domains present in each variant. **b.** Diagram of the *R26^LSL-Cas9-EGFP/+^* (*Cas9-EGFP*) and *Tomato*^T/+^;*Cas9*^T/+^ (*TC*) transgenic mouse models before and after Cre-mediated recombination. Lentiviral Cre induces *Cas9-EGFP* expression in the *Cas9-EGFP* model and *Tomato* and *Cas9* expression in the *TC* model. **c.** Sanger sequencing chromatograms of individual V1 and V3 tumors, showing the expected *Eml4-Alk* rearrangements. **d.** Schematic of tumor initation in *TC* mice by intratracheal injection with Lenti-sgV1/Cre and Lenti-sgV3/Cre vectors. Viral titers per mouse, mouse numbers, and genotypes are indicated. **e.** Representative fluorescent tomato images of tumor-bearing lungs from *TC* mice with V1 or V3 tumors. The scale bars are 5mm. **f.** Lung weights (grams) of *TC* mice with V1 or V3 tumors. Each dot represents a mouse. Unpaired t-test, nominal P-value is shown. **g.** Representative H&E staining of lung sections from *TC* mice with V1 or V3 tumors. The scale bars are 2mm (left) and 100µm (right). **h.** Schematic representation of the cloning strategy. Individual sgRNAs (sgEgfp, sg*Eml4*^V1^, sg*Eml4*^V3^, and sg*Alk*) were cloned individually into U6 donor plasmids. Lenti-sgEgfp-sgV1/Cre and Lenti-sgEgfp-sgV3/Cre were then generated by Gibson assembly, combining U6 sgRNA plasmids with a pLenti/Cre donor plasmid. **i.** Representative H&E and GFP immunohistochemical staining of V3 lung tumor section. Scale bars are 100µm.

**Supplementary Figure 2.**
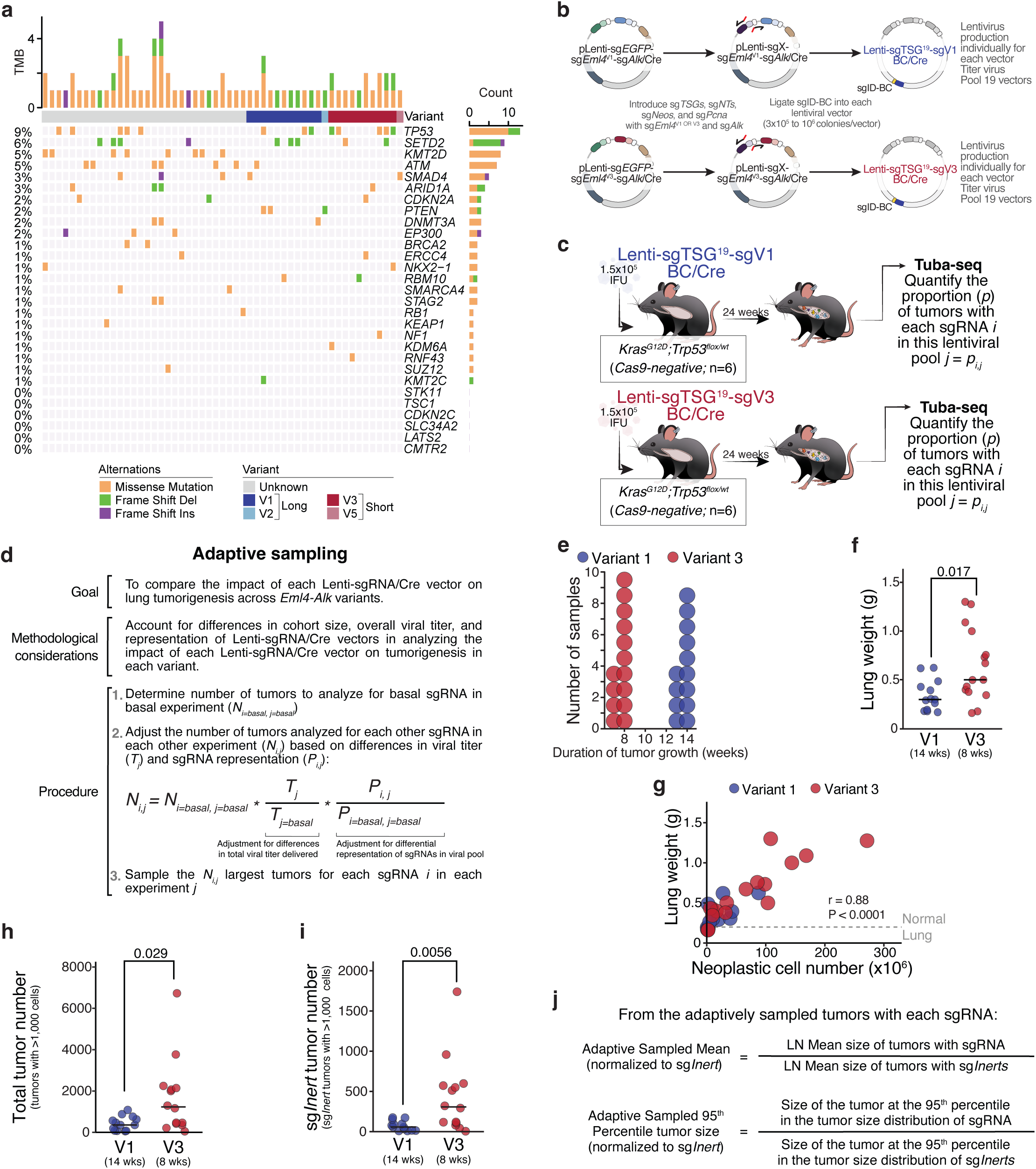
Strategy for quantifying the impacts of tumor suppressor gene inactivation and additional data on V1- and V3-driven tumorigenesis. **a.** Oncoplot showing somatic mutations in genes assayed in this study. Each column represents a specific EML4-ALK sample and the rows represent genes mutated in the samples. Samples are annotated per EML4-ALK variant. Data from GENIE (see methods). **b.** Schematic of diversification of the Lenti-sgTSG^19^-sgV1/Cre and Lenti-sgTSG^19^-sgV3/Cre pools. Each sgRNA targeting a tumor suppressor gene was cloned and individually diversified. **c.** Schematic of tumor initiation in *Kras^G12D^;Trp53^flox/wt^* (*Cas9-negative)* mice with Lenti-sgTSG^19^-sgV1/Cre and Lenti-sgTSG^19^-sgV3/Cre viral pools to determine the representation of Lenti-sgRNA/Cre vectors. **d.** Explanation of adaptive sampling method used in assessing the impact of Lenti-sgRNA/Cre vectors in V1- and V3-driven models (see Methods). **e.** Duration of tumor growth for mice in *Cas9-EGFP* cohorts transduced with Lenti-sgTSG^20^-sgV1/Cre or Lenti-sgTSG^20^-sgV3/Cre viral pools. Each dot is a mouse. **f.** Lung weights of mice in the V1 and V3 cohorts. Each dot represents a mouse and bars indicate median values within each genotype. The P-value was calculated using a two-sided Wilcoxon rank sum test. **g.** Lung weight is highly correlated with the total number of neoplastic cells detected by Tuba-seq for *Cas9-EGFP* mice transduced with Lenti-sgTSG^19^-sgV1/Cre and Lenti-sgTSG^19^-sgV3/Cre. Each dot is a mouse. Pearson correlation coefficient and associated p-value are shown. **h,i.** Total number of tumors larger than 1,000 cells in size (**h**) and number of sg*Inert* tumors larger than 1,000 cells in size (**i**) in the V1 and V3 cohorts. Each dot represents a mouse and bars indicate median values within each genotype. The P-value was calculated using a two-sided Wilcoxon rank sum test. **j.** Schematic of calculation of metrics used to quantify the impacts of tumor suppressor gene inactivation on tumor growth (see Methods).

**Supplementary Figure 3.**
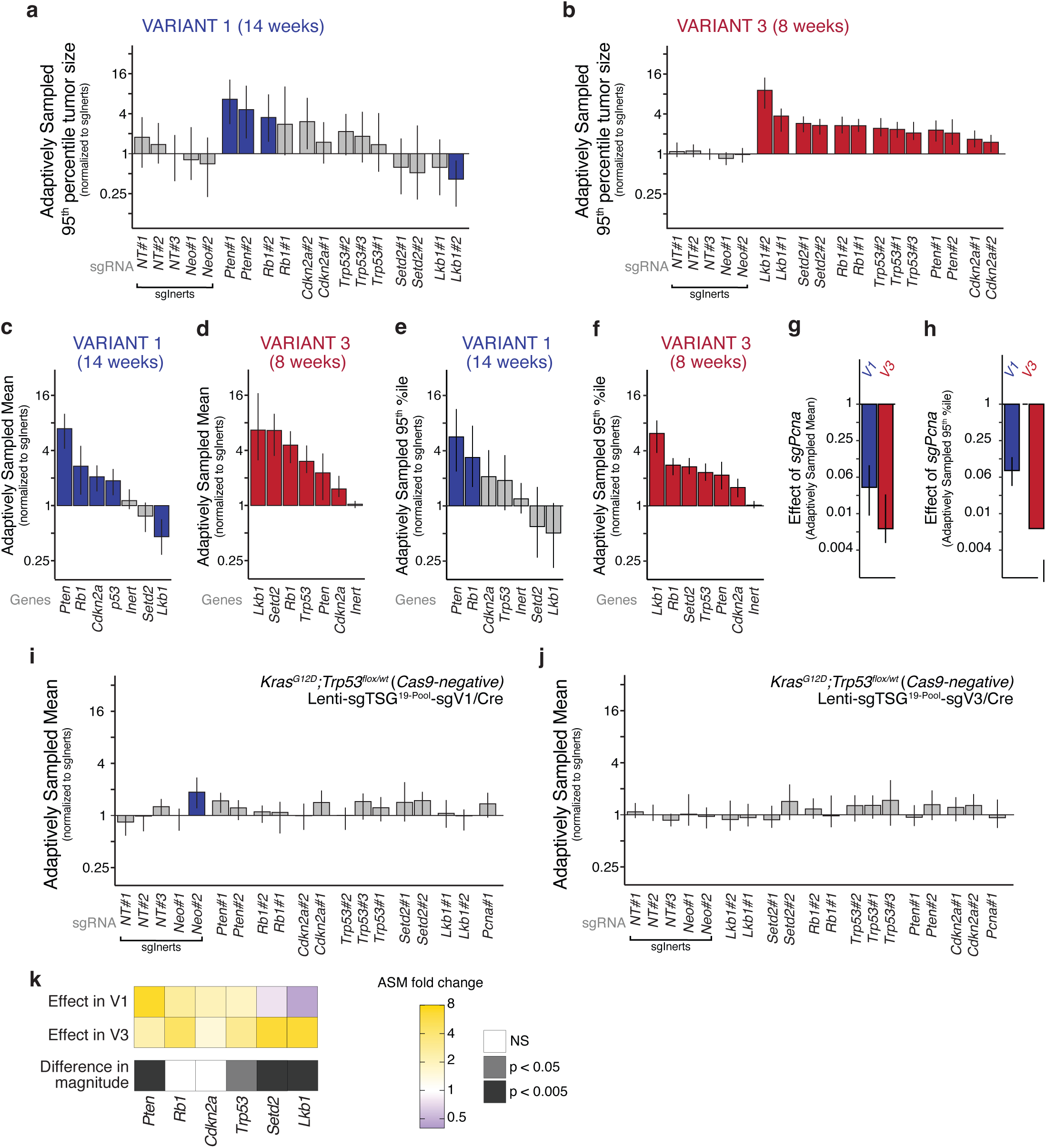
Additional data on the impacts of inactivation of tumor suppressor genes in the Lenti-sgTSG^19^-sgV1/Cre and Lenti-sgTSG^19^-sgV3/Cre. **a-b**. Adaptively sampled 95^th^ percentile tumor size (AS95) of each tumor genotype normalized to the AS95 of sg*Inert* tumors in *Cas9-EGFP* cohorts transduced with the Lenti-sgTSG^19^-sgV1/Cre (**a**, “V1 cohort”) and Lenti-sgTSG^19^-sgV3/Cre (**b**, “V3 cohort”) viral pools. Each bar is a sgRNA. sgRNAs are grouped by gene. sgRNAs that significantly increase or decrease AS95 (two-sided FDR-adjusted P-value) are in color. **c-d**. Adaptively sampled mean tumor size (ASM) relative to sg*Inert* vectors calculated at the gene level in the V1 (**c**) and V3 (**d**) cohorts. Genes that significantly increase or decrease ASM (two-sided FDR-adjusted P-value) are in color. **e-f**. AS95 relative to sg*Inert* vectors calculated at the gene level in the V1 (**e**) and V3 (**f**) cohorts. Genes that significantly increase or decrease AS95 (two-sided FDR-adjusted P-value) are in color. **g-h**. Effect of inactivation of essential gene *Pcna* on tumor growth as measured by ASM (**g**) and AS95 (**h**) relative to sg*Inert* vectors in the V1 and V3 cohorts. **i-j**. ASM relative to sg*Inert* vectors for each sgRNA in the *Kras^G12D^;Trp53^flox/wt^* (*Cas9-negative)* cohorts transduced with the Lenti-sgTSG^19^-sgV1/Cre **(i)** and Lenti-sgTSG^75^-sgV3/Cre **(j)** viral pools. For panels **a**-**j**: Line at Y=1 indicates no impact relative to *sgInert* vectors. P-values and confidence intervals were calculated using nested bootstrap resampling. **k.** Heatmap depicting the tumor suppressive effect of each gene in the V1 and V3 cohorts. Top: impact of inactivating each gene on tumor growth as measured by fold change in ASM relative to control sg*Inert* vectors. The color indicates the magnitude and direction of the effect and genes are ordered by effect size in the V1 cohort. Bottom: significance of the difference in the impact of inactivating each gene in V1- and V3-driven lung tumors.

**Supplementary Figure 4.**
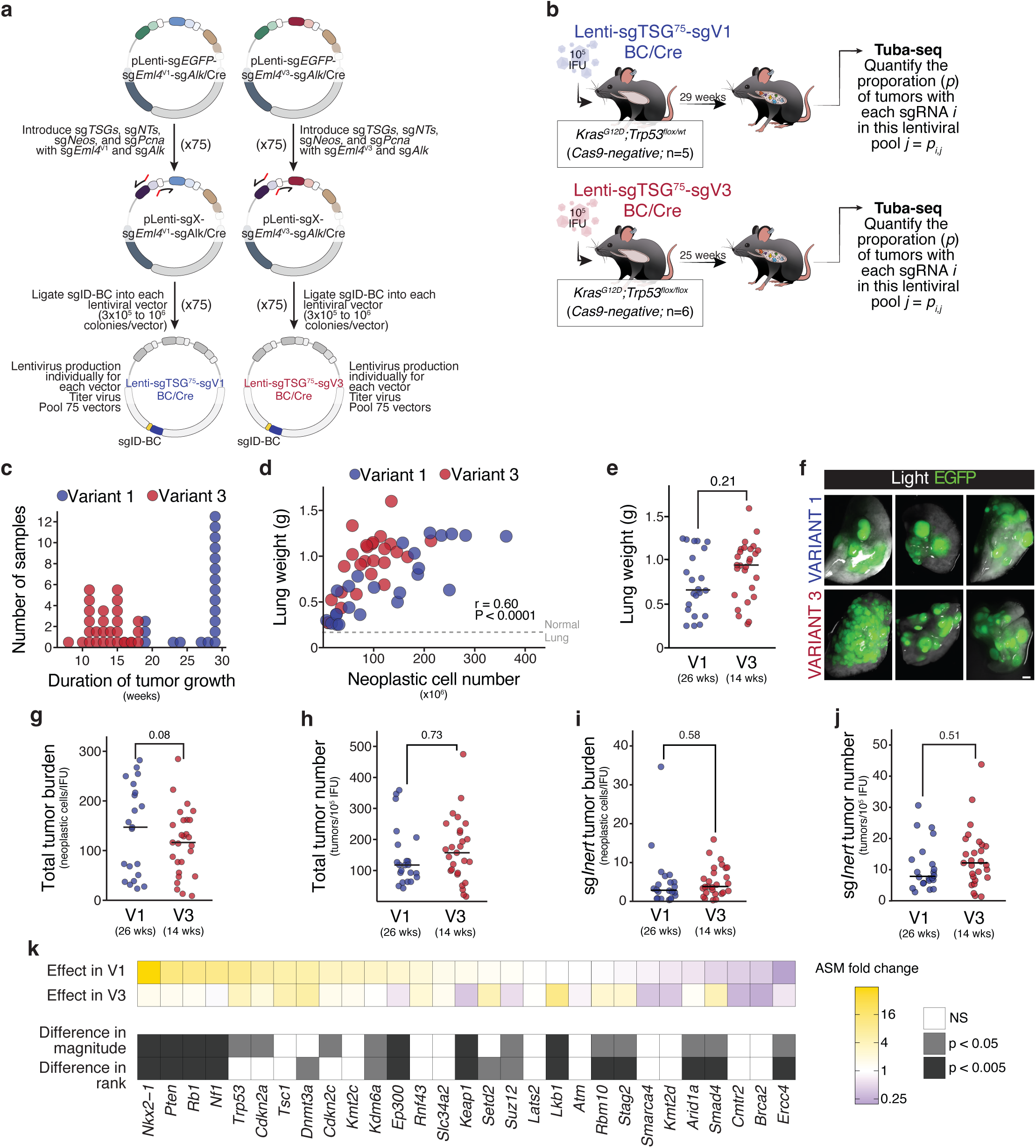
Additional data from broader screen of tumor suppressor function in V1- and V3-driven lung cancer. **a.** Schematic of Lenti-sgTSG^75^-sgV1/Cre and Lenti-sgTSG^75^-sgV3/Cre pools diversification. Each sgRNA targeting a tumor suppressor gene was cloned and individually diversified. **b.** Schematic of tumor initiation in *Cas9-negative* mice with Lenti-sgTSG^75^-sgV1/Cre and Lenti-sgTSG^75^-sgV3/Cre viral pools to determine the representation of Lenti-sgRNA/Cre vectors. **c.** Duration of tumor growth for *R26^LSL-Cas9-EGFP/LSL-Cas9-EGFP^ (Cas9-EGFP)* mice transduced with Lenti-sgTSG^75^-sgV1/Cre (”V1 cohort”) and Lenti-sgTSG^75^-sgV3/Cre (”V3 cohort”). Each dot is a mouse. **d.** Lung weight is highly correlated with the total number of neoplastic cells detected by Tuba-seq for *Cas9-EGFP* mice transduced with Lenti-sgTSG^75^-sgV1/Cre and Lenti-sgTSG^75^-sgV3/Cre. Each dot is a mouse. Pearson correlation coefficient and associated P-value are shown. **e.** Lung weights of mice in the V1 and V3 cohorts. Each dot represents a mouse and bars indicate median values within each cohort. The P-value was calculated using a two-sided Wilcoxon rank sum test. **f.** Fluorescent images of the lungs of *Cas9-EGFP* mice in the V1 and V3 cohort. EGFP fluorescence and bright-field images were merged. Scale bars: 2.5 mm. **g,i.** Total number of neoplastic cells summed across all tumors (**g**) and across all sg*Inert* tumors (**i**) for mice in the V1 and V3 cohorts normalized to the viral titer delivered. Each dot represents a mouse and bars indicate median values within each genotype. The P-value was calculated using a two-sided Wilcoxon rank sum test. **h,j.** Total number of tumors larger than 1,000 cells in size (**h**) and number of sg*Inert* tumors larger than 1,000 cells in size (**j**) for mice in the V1 and V3 cohorts normalized to the viral titer delivered. Each dot represents a mouse and bars indicate median values within each genotype. The P-value was calculated using a two-sided Wilcoxon rank sum test. **k.** Heatmap depicting the tumor suppressive effect of each gene in the V1 and V3 cohorts. Top: impact of inactivating each gene on tumor growth as measured by fold change in the adaptively sampled mean size (ASM) relative to control sg*Inert* vectors. The color indicates both the magnitude and direction of effect and genes are ordered by effect size in the V1 cohort. Bottom: the significance of difference in the impact of inactivating each gene in the V1 and V3 cohorts as measured by both magnitude and rank order of effects.

**Supplementary Figure 5.**
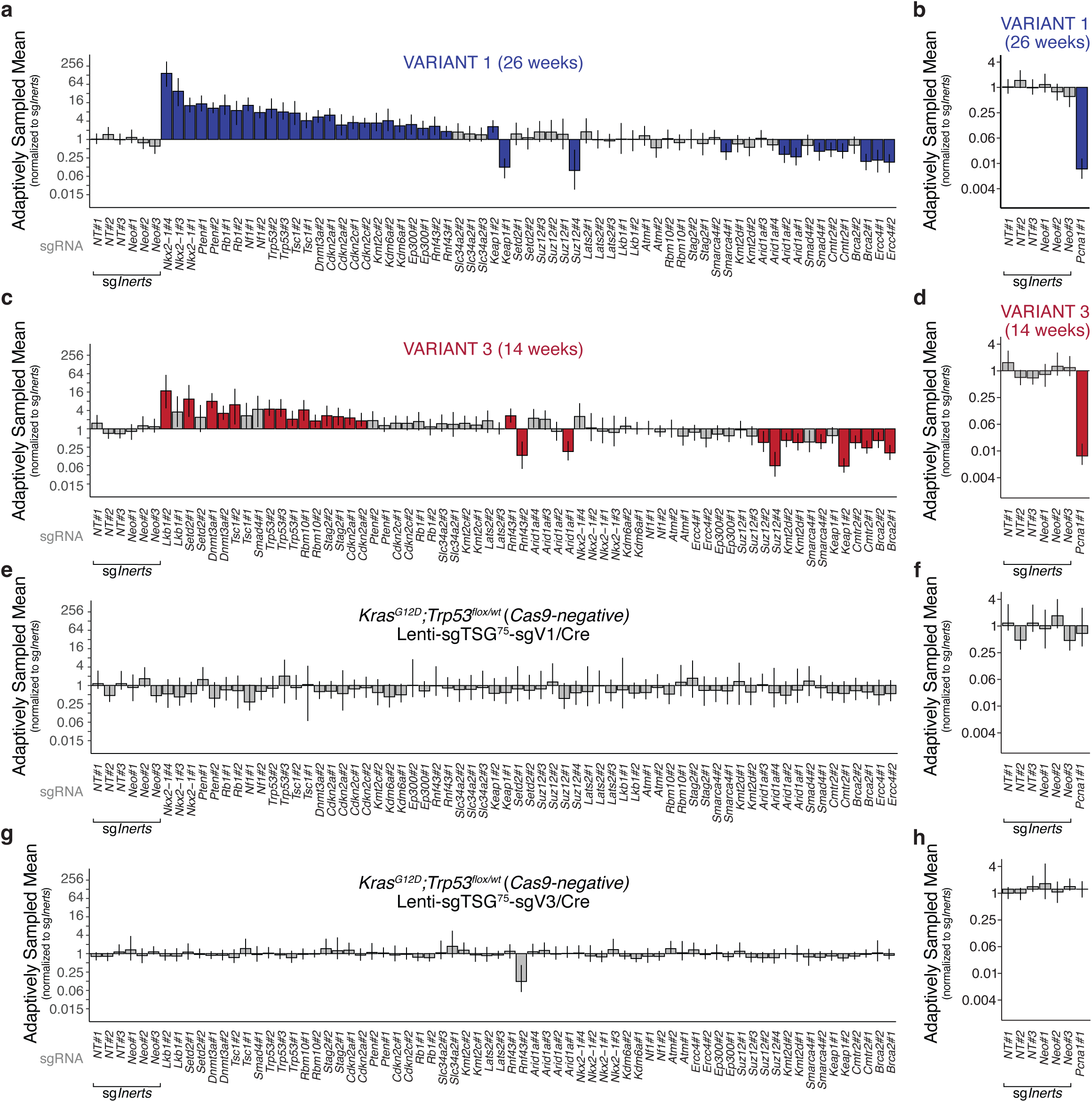
Effects of tumor suppressor gene inactivation at the sgRNA-level in the Lenti-sgTSG75-sgV1/Cre and Lenti-sgTSG75-sgV3/Cre pools. **a, c.** Adaptively Sampled Mean (ASM) tumor sizes relative to sg*Inert* vectors for each sgRNA in the Lenti-sgTSG^75^-sgV1/Cre (**a**) and Lenti-sgTSG^75^-sgV3/Cre (**c**) pools in *Cas9-EGFP* mice. **b, d.** ASM tumor size for *sgPcna* (essential gene control) relative to sg*Inert* vectors in *Cas9-EGFP* cohorts transduced with Lenti-sgTSG^75^-sgV1/Cre (**b**) and Lenti-sgTSG^75^-sgV3/Cre (**c**) pools. **e, g.** ASM tumor sizes relative to sg*Inert* vectors for each sgRNA in theLenti-sgTSG^75^-sgV1/Cre **(e)** and Lenti-sgTSG^75^-sgV3/Cre **(g)** pools in *Cas9-negative* mice. **f, h.** ASM tumor size for *sgPcna* (essential gene control) relative to sg*Inert* vectors in *Cas9-negative* cohorts transduced with Lenti-sgTSG^75^-sgV1/Cre (**f**) and Lenti-sgTSG^75^-sgV3/Cre (**h**) pools. For all panels: ASM is a summary metric of tumor fitness that integrates the impact of inactivating each gene on tumor size and number. Each bar is a sgRNA. sgRNAs that significantly increase or decrease ASM (two-side FDR-adjusted P-value) are in color. P-values and confidence intervals were calculated using nested bootstrap resampling.

**Supplementary Figure 6.**
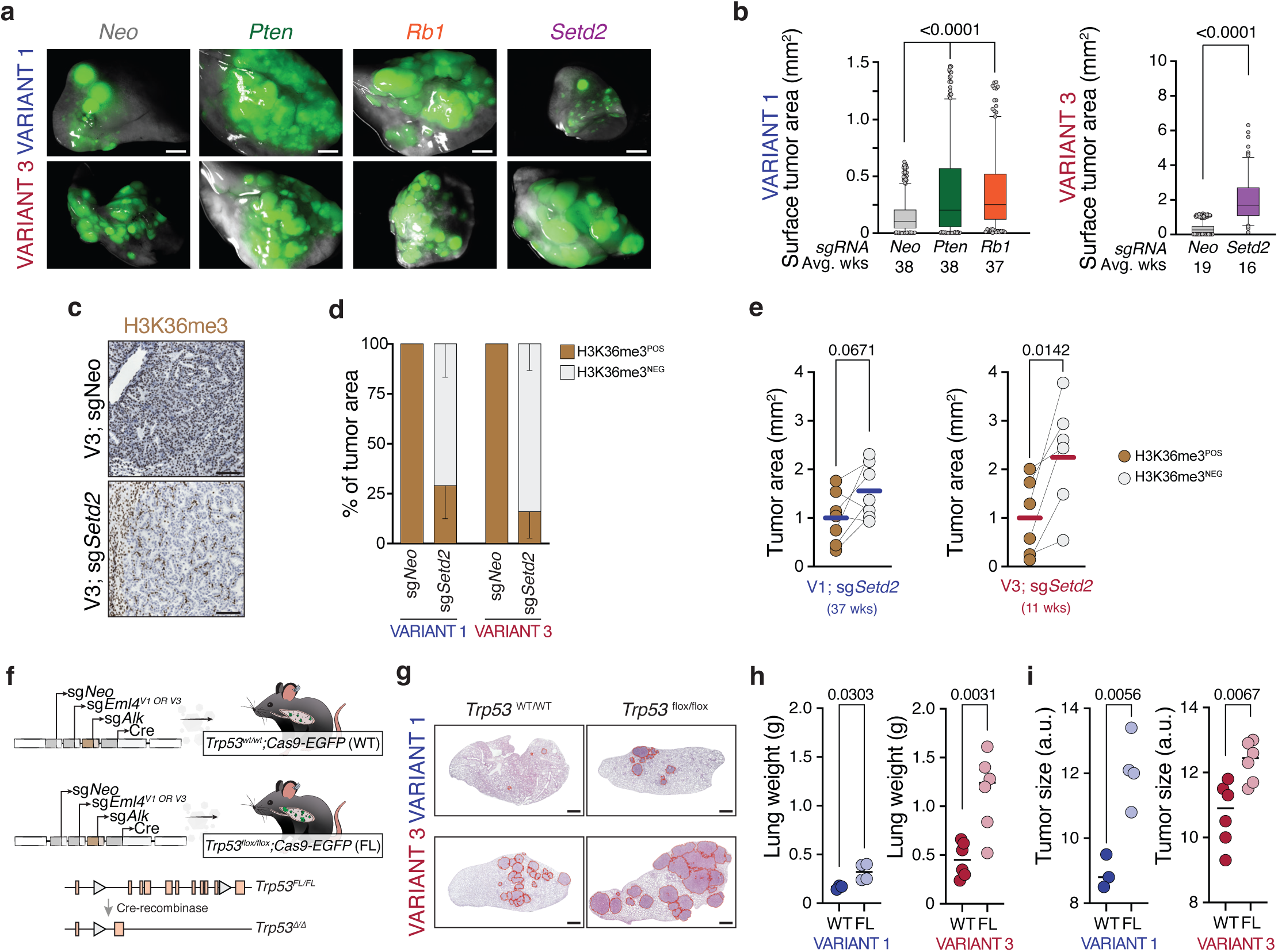
Additional data on validation of EML4-ALK variant-specific effects of tumor suppressor gene inactivation. **a.** Representative light and fluorescent lung images from Lenti-sgTSG-sgV1/Cre and Lenti-sgTSG-sgV3 induced lung tumors. Scale bar is 2 mm. **b.** Surface tumor areas (mm2) from mice with indicated tumor genotypes collected at similar time points. Each point represents a tumor; box-and-whisker plots indicates quartile values (box) and the 5-95 percentile range of tumor sizes (whiskers). Data plotted are a subset of data represented in Figure 4c. Horizontal lines indicate the mean within each group. Two-way ANOVA was performed. Nominal P-values of significant comparison vs. Neo are indicated. **c.** Representative immunohistochemistry of H3K36me3 of a EML4-ALK V3-driven *Setd2*-proficient (top, sg*Neo*) and *Setd2*-deficient (bottom, sg*Setd2*). The same pattern was observed in corresponding EML4-ALK V1-driven tumors. Scale bars are 100 µm. **d.** Percentage of H3K36me3 positive or negative tumor area. EML4-ALK V1 and V3-driven *Setd2*-proficient (sg*Neo*) mice are used as a positive control. **e.** Size (mm^2^) of H3K36me3 positive versus negative tumors from Lenti-sg*Setd2*-sgV1/Cre (left) and Lenti-sg*Setd2*-sgV3/Cre (right) infected mice. Each dot is the mean of tumor size from one mouse. Connecting lines indicate that measurements were taken from the same mouse. Horizontal bars indicate the mean. T-test on paired samples was performed and nominal P-values are shown. **f.** Schematic of tumor initiation in *Cas9-EGFP* and *Trp53^flox/flox^;Cas9-EGFP* mice with Lenti-sg*Neo*-sgV1/Cre and Lenti-sg*Neo*-sgV3/Cre vectors. Mice were injected with 1.5x10^6^ IFU/mouse. **g.** Representative H&E staining of lung sections from *Cas9-EGFP* and *Trp53^flox/flox^;Cas9-EGFP* mice with V1 and V3 lung tumors. Single tumors are indicated in red. Scale bars are 1 mm. **h.** Lung weight (grams) from V1 and V3 *Cas9-EGFP* (WT) and *Trp53^flox/flox^;Cas9-EGFP* (FL) mice. Each dot represents a mouse. Unpaired t-test, nominal P-value is shown. **i.** Tumor sizes from V1 and V3 *Cas9-EGFP* (WT) and *Trp53^flox/flox^;Cas9-EGFP* (FL) mice. Each dot represents the median of tumor size per mouse. Unpaired t-test, nominal P-value is shown.

**Supplementary Figure 7.**
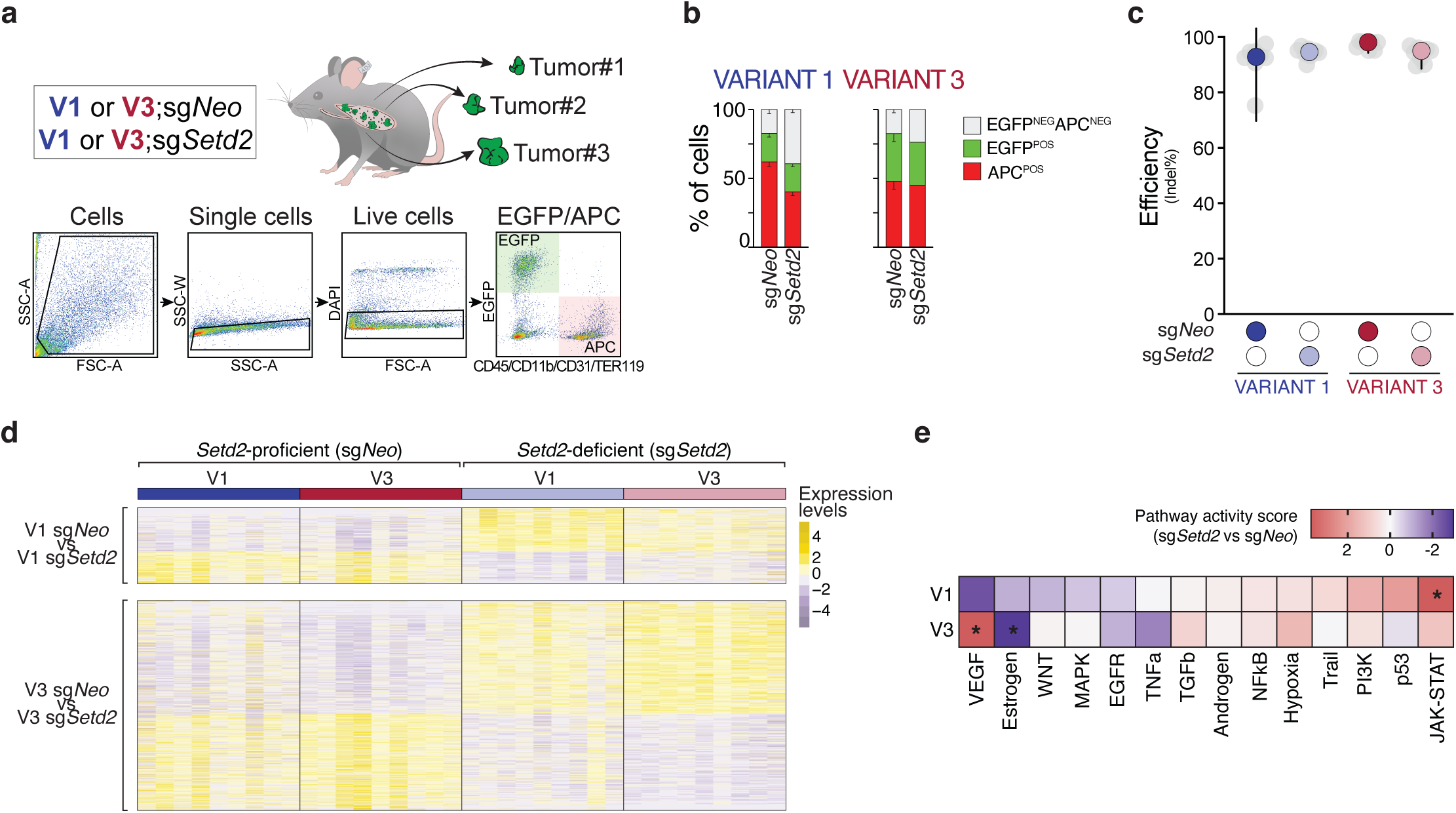
Generation of Setd2-proficient and -deficient V1- and V3-driven cancer cells for transcriptomic analysis. **a.** Schematic of the experimental workflow to isolate Setd2-proficient and Setd2-deficient V1- and V3-driven cancer cells. Single tumors from each genotype were processed for fluorescence-activated cell sorting (FACS; EGFP+/CD45-/CD11b-/CD31-/TER119-). **b.** Frequency of EGFP (green), CD45/CD11b/CD31/TER119 (APC, red) and double negative cells from (a). **c.** Percentage of insertion-deletion (indel) from sorted neoplastic cells. Each colored point is the mean of indels for that genotype. Grey points indicate individual tumors. Vertical lines indicate the coefficient of interval. **d.** Heatmaps showing Z-scored normalized counts for differentially expressed genes from indicated contrasts across all *Setd2*-deficient and *Setd2*-proficient V1- and V3-driven tumors. **e.** Inferred pathway activity scores comparing *Setd2*-deficient (sg*Setd2*) to *Setd2*-proficient (sg*Neo*) V1- and V3-driven tumors. The inferred activity is based on the t-values of the differentially expressed genes for each comparison. Positive scores (red) indicate that a pathway is more active with *Setd2* inactiavtion; negative scores (blue) indicate that a pathway is less active with *Setd2* inactivation. Stars denote that a pathway is differentially activated upon *Setd2*-inactivation (p<0.05, PROGENy multivariate linear model).

**Supplementary Figure 8.**
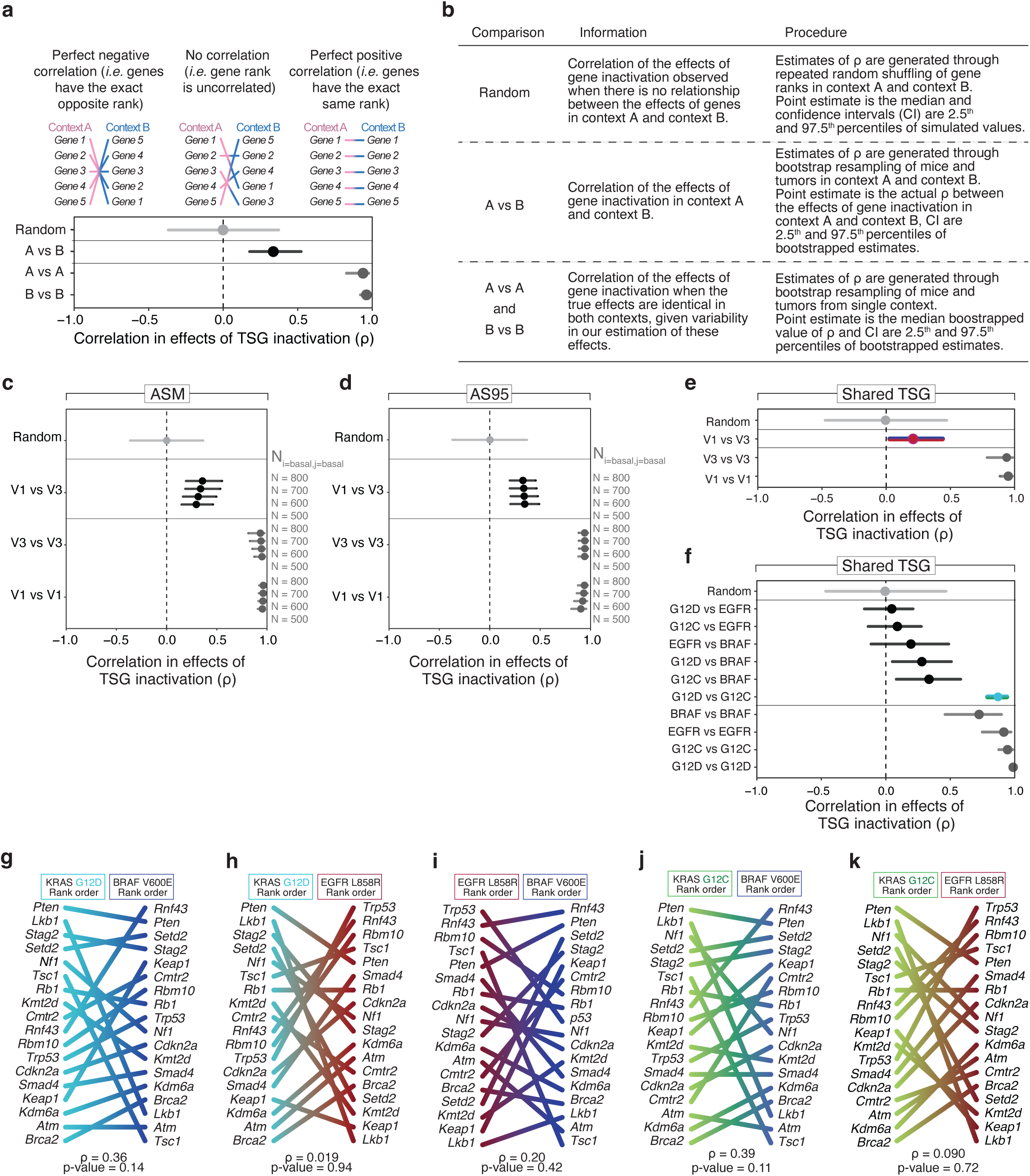
Differences in tumor suppressor function between EML4-ALK V1 and V3 variants are robust to analysis choices and gene sets. **a-b**. Example plot of correlation analysis contrasting two oncogenic contexts (**a**) and accompanying explanation of comparisons (**b**). **c.** Correlation in the rank order of tumor suppressive effects in V3 and V1 mice calculated across various values of Ni=basal,j=basal. Genes were ranked by fold change in ASM relative to sg*Inert* vectors within each analysis. Values of ρ corresponding to Ni=basal,j=basal=700 are also shown in Figure 5a. **d.** Correlation in the rank order of tumor suppressive effects in V3 and V1 mice calculated across various values of Ni=basal,j=basal with genes ranked by fold change in adaptively sampled 95^th^ percentile tumor size relative to sg*Inert* vectors within each analysis. **e.** Correlation in the rank order of tumor suppressive effects in the V1 and V3 cohorts for the subset of genes that overlap with Blair *et al*. Genes were ranked by fold change in ASM relative to sg*Inert* vectors. **f.** Correlation in the rank order of tumor suppressive effects for the subset of genes that overlap with this study between KRAS G12D, KRAS G12C, BRAF, and EGFR mice from Blair *et al*. Genes were ranked by fold change in 95^th^ percentile tumor size relative to sg*Inert* vectors. **g-k.** Comparisons of the rank order of tumor suppressive effects between indicated pairs of oncogenes and oncogene variants using data from Blair *et al*. Genes were ranked by fold change in 95^th^ percentile tumor size relative to sg*Inerts* using an adaptive cut-off. Only genes assayed in both the Lenti-[sgTSG-sgV1/V3]^Pool^BC/Cre pools and in Blair *et al*. are shown.

**Supplementary Figure 9.**
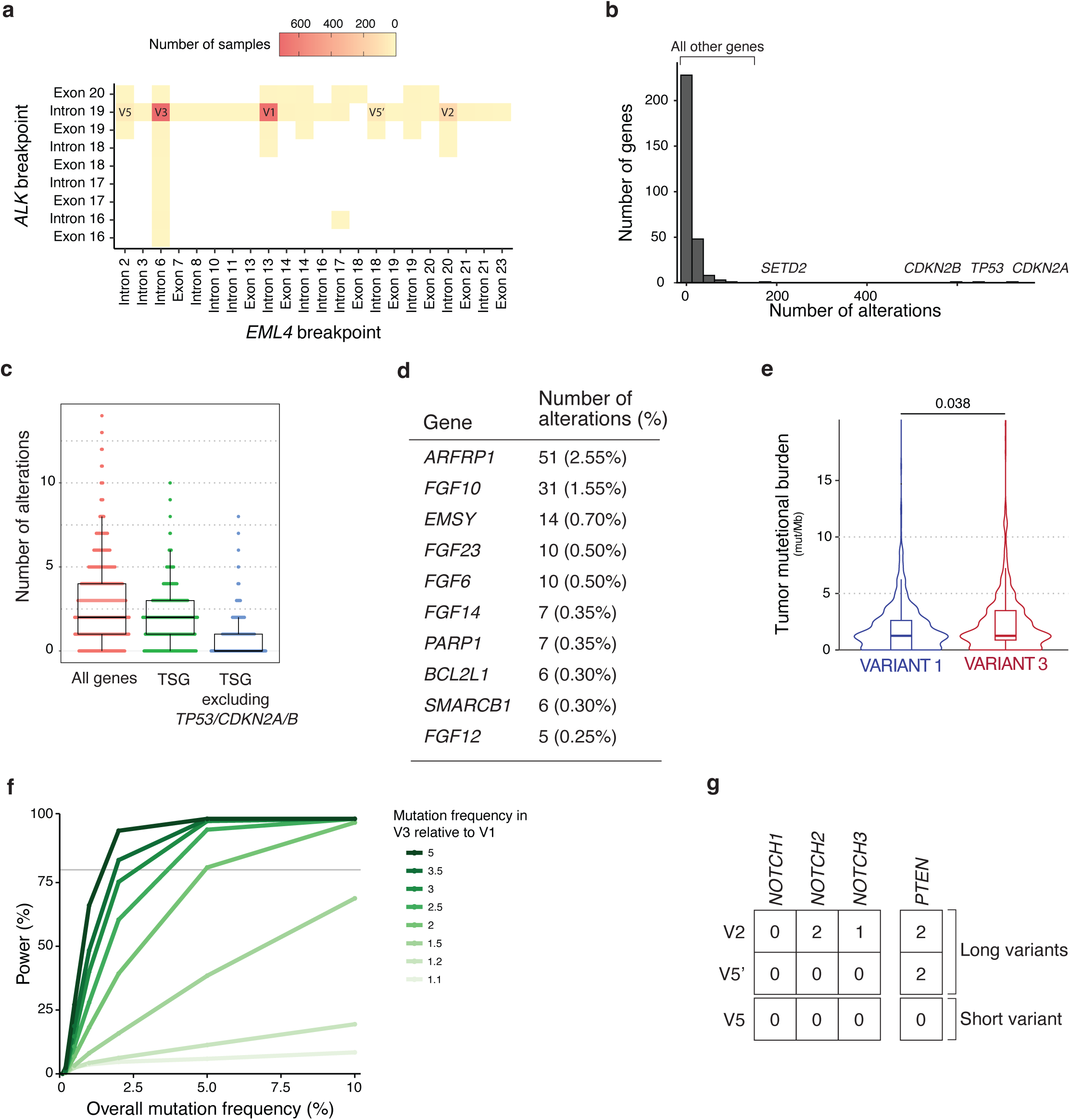
Additional data on the genomic landscape of EML4-ALK-driven lung cancer. **a.** Breakpoints in *EML4* and *ALK* for all samples in the EML4-ALK non-small cell lung cancer (NSCLC) cohort. **b.** Distribution of alteration counts for all genes queried in Foundation Medicine’s sequencing panel. Genes that are commonly altered in the EML4-ALK NSCLC cohort are indicated (*CDKN2A*, *TP53*, *CDKN2B*, *SETD2*); all other genes are altered at frequencies below 5%. **c.** Number of alterations (excluding EML4-ALK fusions) in samples in the EML4-ALK NSCLC cohort. Each point is a sample. **d.** Genes with recurrent alterations in the EML4-ALK NSCLC cohort not previously shown to be altered in EML4-ALK-driven lung cancer. Only genes altered in at least 5 samples are shown. **e.** Tumor mutational burden (TMB) is similar but slightly higher in tumors with EML4-ALK V3 relative to tumors with EML4-ALK V1. Wilcoxon test p-value is reported. **f.** Power to detect differences in mutation frequency between V1- and V3-driven tumors given the size of the V1 and V3 cohorts, overall mutation frequency, and varying relative frequencies of mutation in V1-driven versus V3-driven samples. For example, for a gene altered at an overall frequency of 2% (a prevalence higher than roughly 95% of the genes assayed), mutations would need to occur more than three times as frequently in V3 samples than V1 samples for an interaction to be detected with 80% power. Power was estimated by simulation assuming a binomial distribution for mutation counts in each cohort.10,000 simulations were performed for each scenario. Gray line indicates 80% power. **g.** Alteration counts for *NOTCH* genes and *PTEN* in indicated “long” and “short” rare EML4-ALK variants.

